# Targeting tGLI1, a novel mediator of tumor therapeutic resistance, using Ketoconazole sensitizes glioblastoma to CDK4/6 therapy and chemoradiation

**DOI:** 10.1101/2025.02.20.639359

**Authors:** Yang Yu, Austin Arrigo, Ankush Chandra, Chuling Zhuang, Mariana K. Najjar, Munazza S. Khan, Dongqin Zhu, Antonio Dono, Roy E. Strowd, Nitin Tandon, Jay-Jiguang Zhu, Sigmund H. Hsu, Yoshua Esquenazi, Michael Chan, Hui-Wen Lo

## Abstract

Glioblastoma (GBM) remains the most aggressive primary brain tumor in adults, with no effective treatments. While cyclin-dependent kinase 4/6 inhibitors (CDK4/6is) show clinical promise in some cancers, they have not significantly improved survival in GBM patients. This lack of response is attributed to the treatment-resistant glioma stem cell (GSC) population. We previously identified truncated glioma-associated oncogene homolog 1 (tGLI1) as a novel transcription factor promoting GSCs; however, its role in CDK4/6i resistance has never been investigated in any cancer type. Here, we found positive correlations between tGLI1 and CDK4/6 therapeutic resistance in patient datasets and *in vitro* studies. Pharmacological inhibition of tGLI1 using FDA-approved ketoconazole (KCZ), a tGLI1-specific inhibitor, sensitized GBM and GSCs to CDK4/6is. KCZ+CDK4/6i combination therapy demonstrated synergistic anti-proliferative effects, significantly inhibiting GBM stemness and cell cycle progression while increasing apoptosis. The combination was more efficacious than monotherapies in two orthotopic GBM mouse models. tGLI1 promoted GBM resistance to radiation therapy and temozolomide, while KCZ potentiated effects of these treatments. Collectively, we report for the first time that tGLI1 is a novel mediator of GBM resistance to CDK4/6is, and KCZ sensitizes GBM to CDK4/6is, thereby supporting future clinical utility of novel KCZ+CDK4/6i combinatorial therapy for GBM patients.

## Introduction

Glioblastoma (GBM) is the most common primary malignant brain tumor diagnosed in adults, comprising about half of all malignant brain tumors (1, 2). Despite advancements in tumor biology and novel therapies, the standard-of-care therapy for GBM patients remains maximal safe tumor resection followed by radiation and chemotherapy (3), with an abysmal median overall survival of ∼14 months and less than 7% five-year survival rate (4). Several factors including tumor heterogeneity, high proliferative rate, invasiveness, blood-brain-barrier (BBB), and blood-tumor-barrier (BTB) contribute to making the effective treatment of GBM a major challenge (5). Targeted therapies have shown promise in many cancer types with increased treatment efficacy and safety compared to traditional chemotherapies (6). However, targeted therapies for GBM are limited, with bevacizumab being the only FDA-approved targeted therapy and having no benefit on patient overall survival (OS) (7, 8). Together, there is an urgent unmet need to identify new drug targets and new effective therapies for GBM patients.

Alteration in the Cyclin-dependent kinase 4/6 (CDK4/6) cell cycle pathway are common across many cancers such as breast cancer and melanoma and have been studied as targets for novel therapies (9). The high frequency of CDK4/6 pathway alterations has led to the development of multiple CDK4/6 specific inhibitors, including abemaciclib, palbociclib, and ribociclib (10). Moreover, the use of CDK4/6 inhibitors (CDK4/6is) has shown promise across multiple cancer types including lung, melanoma, colorectal, and breast cancer (11-14). Excitingly, these inhibitors have recently been approved by the U.S. Food and Drug Administration (FDA) for the treatment of hormone receptor-positive, HER2-negative breast cancer (15).

The CDK4/6 pathway has also been found to play an important role in GBM. Specifically, CDK4 overexpression has been shown to promote GBM cell growth and is correlated with worse patient survival (16). CDK4/6 have also been shown to promote a more aggressive GBM phenotype, the mesenchymal subtype, by promoting resistance to cell cycle arrest, apoptosis, and radiation resistance (17-19). While CDK4/6is have shown promising results in treating GBM *in vitro* and *in vivo* (14, 20), multiple clinical trials assessing the use of CDK4/6 monotherapies in GBM patients have shown limited efficacy (13, 21, 22). This discrepancy between the data from preclinical studies and outcomes of clinical trials strongly suggests CDK4/6i resistance as a potential mechanism driving limited efficacy.

Truncated glioma-associated oncogene homolog 1 (tGLI1) was first discovered by our group as a novel splice variant of the zinc-finger transcription factor GLI1 with a gain-of-function downstream effect on the SHH-PTCH1-SMO axis, that has tumor-specific expression in both breast cancer and GBM (23-25). tGLI1 drives the mesenchymal subtype of GBM, and promotes angiogenesis and invasion through upregulation of *CD24, VEGF-C, TEM7*, and *heparinase*, as well as, enrichment of the glioma stem cell (GSC) population through upregulating *CD44* (23, 26-28). Recently, we identified a tGLI1 specific inhibitor and an FDA-approved antifungal, Ketoconazole (KCZ), that specifically inhibits tGLI1-expressing breast cancer stem cells and decreases breast cancer brain metastatic burden in mouse models (29). However, KCZ-mediated inhibition of tGLI1 in GBM has not been investigated.

Given the GSC population plays a critical role in GBM tumor maintenance, proliferation, treatment resistance, and recurrence (30), targeting this cell population may aid in improving treatment efficacy against GBM. Thus, in this study, we sought to characterize the role of tGLI1 in promoting CDK4/6i resistance and study the effects of KCZ+CDK4/6i combination targeted therapy against GBM using *in vitro* experiments and two orthotopic GBM xenograft models. Our data showed, for the first time, that tGLI1 renders GBM cell and GSCs more resistant to CDK4/6i treatment, tGLI1 specific knockdown potentiates the cells to CDK4/6i, and the novel KCZ+CDK4/6i combination treatment synergistically inhibited GSC formation, induced cell cycle arrest, increased apoptosis, decreased tumor growth, and reduced angiogenesis *in vitro* and *in vivo*. Furthermore, we found that tGLI1 overexpression renders GSCs more resistant to radiation (RT) and temozolomide (TMZ), and conversely, KCZ sensitizes tGLI1-expressing GSCs to radiation and TMZ treatments. Together, these findings reveal tGLI1 as a promising pharmacological target to sensitize GBM to standard-of-care as well as CDK4/6-targeted therapy.

## RESULTS

### CDK4/6 pathway is altered in GBMs

To determine the frequency of CDK4/6 pathway alteration in GBM patients, we performed bioinformatic analyses using cBioPortal (31-33). After excluding isocitrate dehydrogenase mutant (IDH-mt) patients, in accordance with the current diagnostic criteria for GBM, we found that roughly 70% of GBM patients had some genetic alteration of key CDK4/6 pathway components (**Fig. 1A**). Interestingly, CDKN2A and CDKN2B deletions were the most frequent alterations in patients at 55% and 53% respectively, with the majority of patients having loss of both genes (**Fig. 1A**). Furthermore, we found alterations in other key CDK4/6 pathway components (CDK4, CDK6, RB1), to be mostly exclusive to CDKN2A/B deletion in patients (**Fig. 1A**). Patients with any CDK4/6 pathway alteration had worse overall survival (Median: 13.76 months vs 17.18 months, p<0.0001) and disease-free survival (DFS) (Median: 7.10 months vs 9.56 months, p=0.0008) compared to patients without any CDK4/6 pathway alterations (**Fig. 1B-C**). While no significant difference was observed in OS or DFS in patients with only CDK4/6 amplification versus unaltered patients, OS trended to be worse in patients with CDK4/6 amplification (**Supp Fig. 1 and 2**). Additional patient samples from the Wake Forest Brain Tumor Tissue Bank were analyzed (N=33) which also showed that CDKN2A/B loss and CDK4/6 amplification are frequent in GBM patients (**Fig. 1D-E**). Furthermore, CDK4/6 amplifications are found in both newly diagnosed and recurrent patient samples (**Fig. 1E**). Similarly, patient samples collected at the University of Texas Health Science Center at Houston (N=281) showed similar results (**Supp. Fig. 3**), with over two-thirds of primary and recurrent patients having alterations in the CDK4/6 pathway (**Supp. Fig. 3A-B, Supp Fig. 4A**). Additionally, in matched primary and recurrent samples, 6% of patients showed a gained CDK4/6 pathway mutation in their recurrent sample versus primary tumor (**Supp. Fig. 4B**). Together, these findings demonstrate the critical role of CDK4/6 pathway alterations in driving primary and recurrent GBM and the importance of inhibiting this pathway.

**Figure 1:**
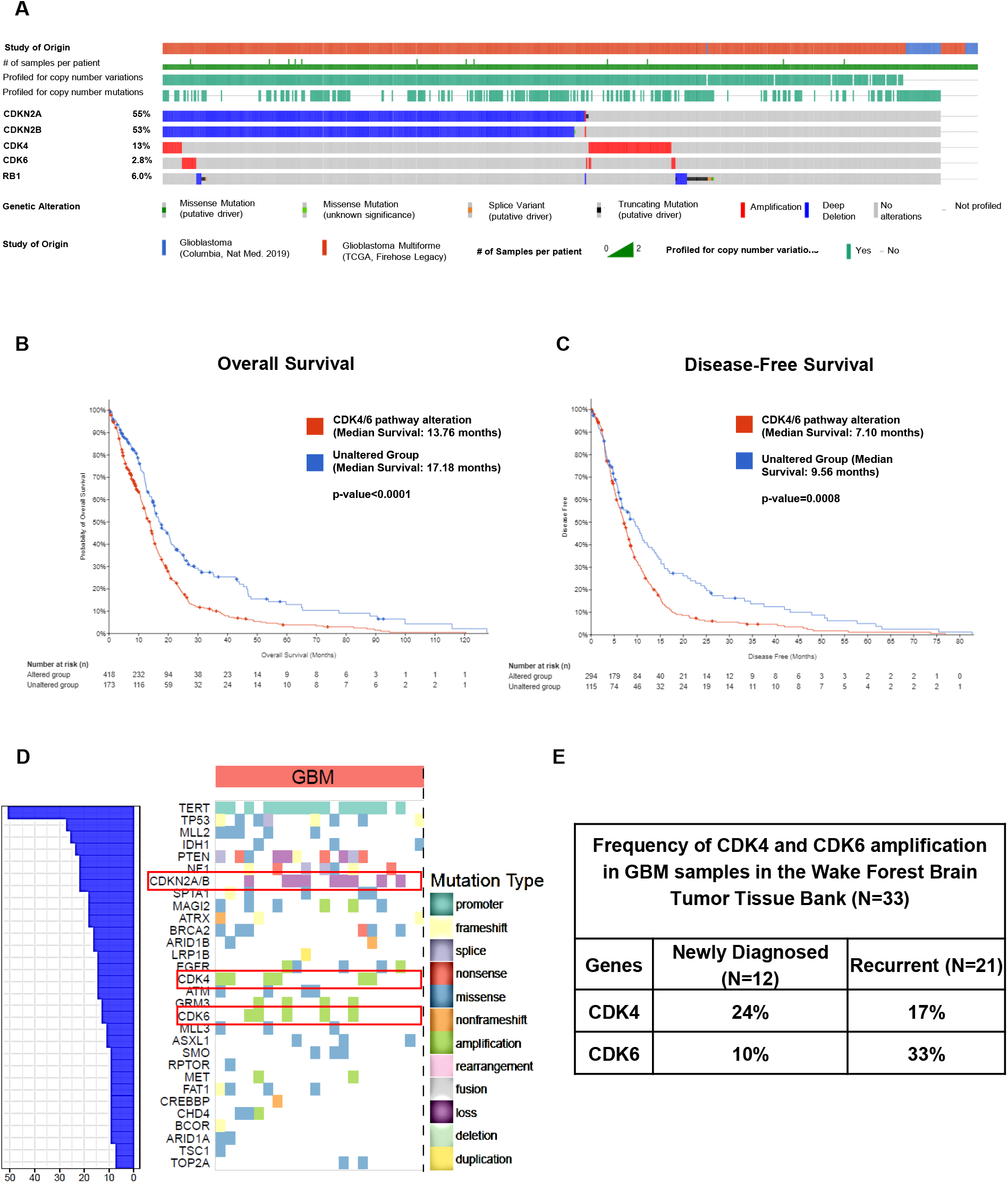
CDK4/6 pathway is altered in the majority of GBM patients. **(A)** CDK4/6 pathway alterations are common in GBM patients. Oncoprint of CDK4/6 pathway genomic alterations in GBM (IDH-WT) patients generated from cBioPortal GBM datasets (N=642). **(B-C)** Patients with any CDK4/6 pathway alteration have worse OS and DFS compared to the unaltered group. Kaplan-Meier analysis of GBM (IDH-WT) patients’ OS and DFS with or without CDK4/6 pathway alterations from cBioPortal GBM datasets (N=642). **(D)** GBM patient samples (newly diagnosed and recurrent) collected at Wake Forest Comprehensive Cancer Center (WFCCC) (N=33) frequently have genomic alterations in CDKN2A/B, CDK4, or CDK6. **(E)** WFCCC patients have 24% CDK4 and 10% CDK6 amplification in newly diagnosed patients (N=12), and 17% CDK4 and 33% CDK6 amplification in recurrent samples (N=21). Log rank test was used to compute p-values.

### tGLI1 pathway activation is correlated with CDK4/6i resistance in GBM

Due to the lack of response to CDK4/6is as observed in multiple clinical trials, we next sought to identify potential mechanisms underlying this therapeutic resistance (21, 22). Using a published CDK4/6i resistance signature (CIRS) and Gene Set Enrichment Analysis (GSEA) (26, 27), we found three cancer signaling pathways to be significantly enriched in patients with a high CIRS score (**Fig. 2A**), namely β-catenin, Sonic HedgeHog (SHH), and PI3K (**Fig. 2B-D**). Since it is well-established that the SHH pathway promotes GSC population, which is the known driver of treatment resistance in GBM(34), we decided to focus on investigating the SHH pathway and its potential role in CDK4/6i resistance. Given that our recent work has shown glioma GLI1 and tGLI1 as the downstream effectors of the SHH pathway (23), we next explored GLI1 (GLI1 activation signature) and tGLI1 activity (tGLI1 activation signature—tGAS) in relation to CIRS score in GBM patients. By performing Pearson correlation analysis, we found tGLI1 activity to be significantly and positively correlated with CIRS in GBM patients, with no correlation between GLI1 and CIRS (**Fig. 2E-F**). Furthermore, patients with mesenchymal GBM (mGBM) were found to have a high CIRS/tGAS combined signature score as compared to other GBM subtypes (**Fig. 2G**). mGBM exhibits increased stemness, invasion, and treatment resistance, further suggesting the role of tGLI1-mediated GBM stemness as a potential mechanism of CDK4/6i resistance (35).

**Figure 2:**
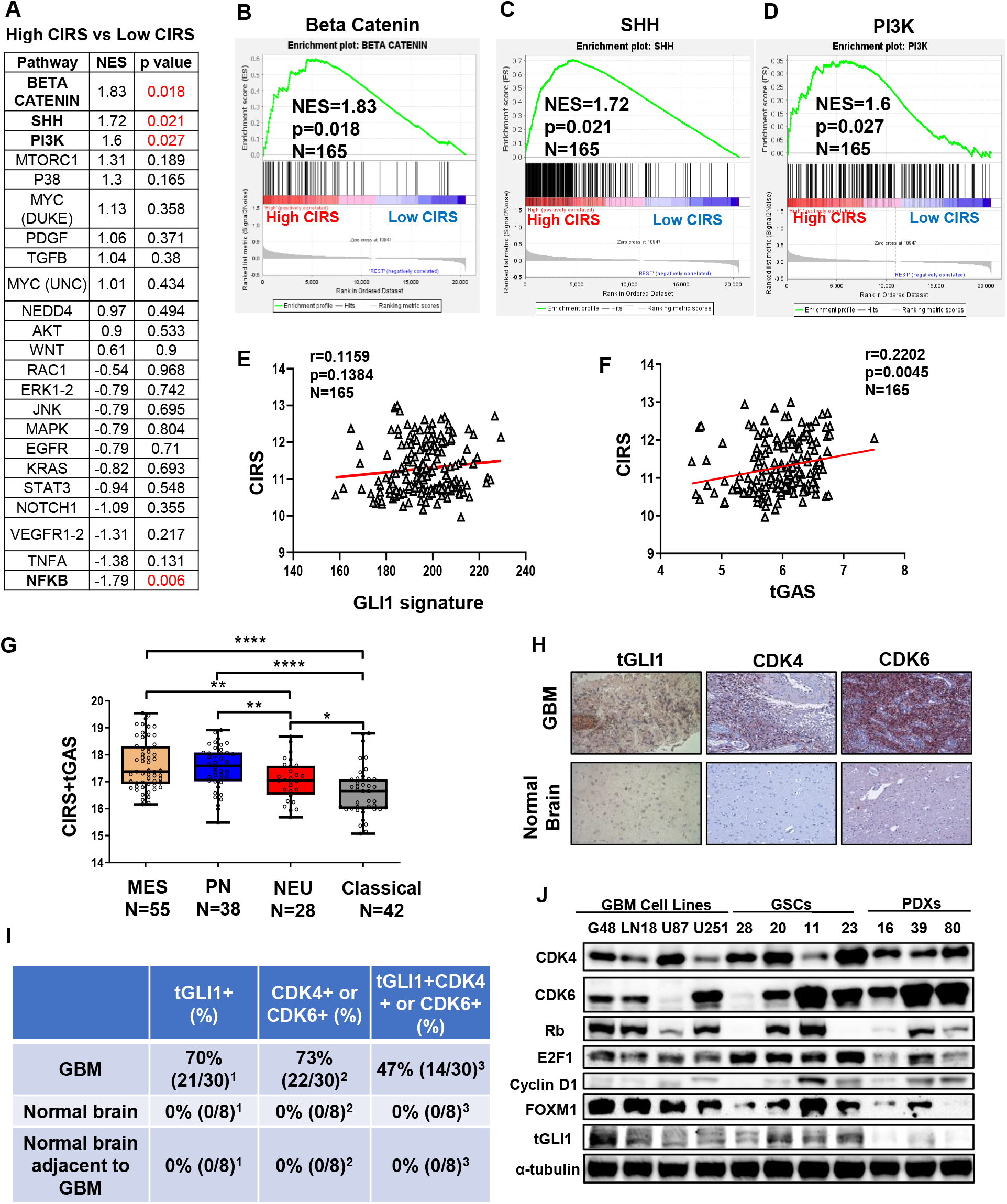
tGLI1 pathway activation is positively correlated with CDK4/6 inhibitor resistance. **(A)** Signaling pathways were analyzed for enrichment in High vs Low CIRS patients, with their respective Normalized Enrichment Scores (NES) and p-values. β-catenin, SHH, and PI3K pathways were the three most enriched. **(B-D)** Representative GSEA plots for β-Catenin (B), SHH (C), and PI3K (D). **(E-F)** tGAS, not GLI1 signature, is significantly and positively correlated with CIRS in GBM patients. Pearson correlation plot of CIRS versus GLI1 signature (E) or tGAS (F). **(G)** Combined CIRS/tGAS signature is highly expressed in the mesenchymal subtype of GBM patients from the GSE4290 dataset. **(H)** Representative images of tissue microarray (GL2083) co-stained for tGLI1, CDK4, or CDK6, consisting of 30 GBM samples, 8 normal brain tissues, and 8 cancer adjacent normal tissues. **(I)** Corresponding values of positive expression from stained GL2083 slides showing high expression of tGLI1, CKD4/6, and tGLI1 and CDK4/6 in GBM patient samples. Fisher’s exact test was used to compute p-values (^1^p=0.0005, ^2^p=0.0003, and ^3^p=0.0168 between GBM and normal/adjacent brain samples) **(J)** Western blot panel containing GBM cell lines, GSCs, and PDXs samples were analyzed for levels of CDK4/6 pathway proteins and tGLI1. Student’s t-test was used to compute p-values. *p<0.05, **p<0.01, ***p<0.001, ****p<0.0001.

Given the correlation between tGLI1 activity with CIRS in publicly available datasets, we next determined whether this correlation remained true in GBM patient specimens and cell lines. A tissue microarray (GL2083) containing 30 GBMs, 8 normal brain, and 8 GBM adjacent tissue samples was stained for tGLI1 and CDK4/6 proteins using immunohistochemistry (IHC) (representative images in **Fig. 2H**). Results revealed that both tGLI1 and CDK4 or CDK6 proteins were highly expressed in GBM patients (70% and 73%, respectively) with no signal in normal or tumor adjacent tissues (**Fig. 2H-I**). Additionally, 47% of patients showed co-expression of tGLI1 and CDK4/6 proteins in GBM (**Fig. 2H-I**). Western blot analysis using multiple GBM cell lines, GSCs, and patient derived xenograft (PDX) samples revealed that tGLI1 is co-expressed with CDK4/6 in the majority of the GBM samples tested (**Fig. 2J**). Additionally, E2F Transcription Factor 1 (E2F1) and Forkhead Box M1 (FOXM1), two downstream targets of the CDK4/6 pathway, were also highly expressed in tGLI1 positive GBM lines (**Fig. 2J**). Of note, loss of Rb and CDK6 was observed in some of the cell lines, which have been shown to play an important role in promoting CDK4/6i resistance. Thus, G48 (CDK6+ and Rb+) and GSC28 (CDK6-and Rb-) lines were selected as representative cell lines for downstream analysis (**Fig. 2J**) (36). Together, these data show that tGLI1 and CDK4/6 pathway proteins are co-expressed in GBM, suggesting that tGLI1 activity is correlated with the observed CDK4/6i resistance in patients.

### tGLI1 promotes GBM resistance to CDK4/6i in vitro

The GSC population is one of the primary drivers of GBM treatment resistance. Therefore, we next investigated the role of tGLI1 in promoting CDK4/6i resistance in the GSC population. Neurosphere formation assays, to enrich the stem cell population, were performed using G48 and GSC-28 cell lines stably overexpressing a vector, GLI1, or tGLI1 with CDK4/6i treatment (abemaciclib or palbociclib) (**Fig. 3A-E**). tGLI1-overexpressing G48-tGLI1 and GSC-28-tGLI1 lines demonstrated greater resistance to abemaciclib treatment as compared to their vector and GLI1 controls (**Fig. 3A-B; Supp. Fig.6A and 6B**). Similar results were found with Palbociclib treatment (**Fig. 3C-D; Supp. Fig. 6C and 6D**).

**Figure 3:**
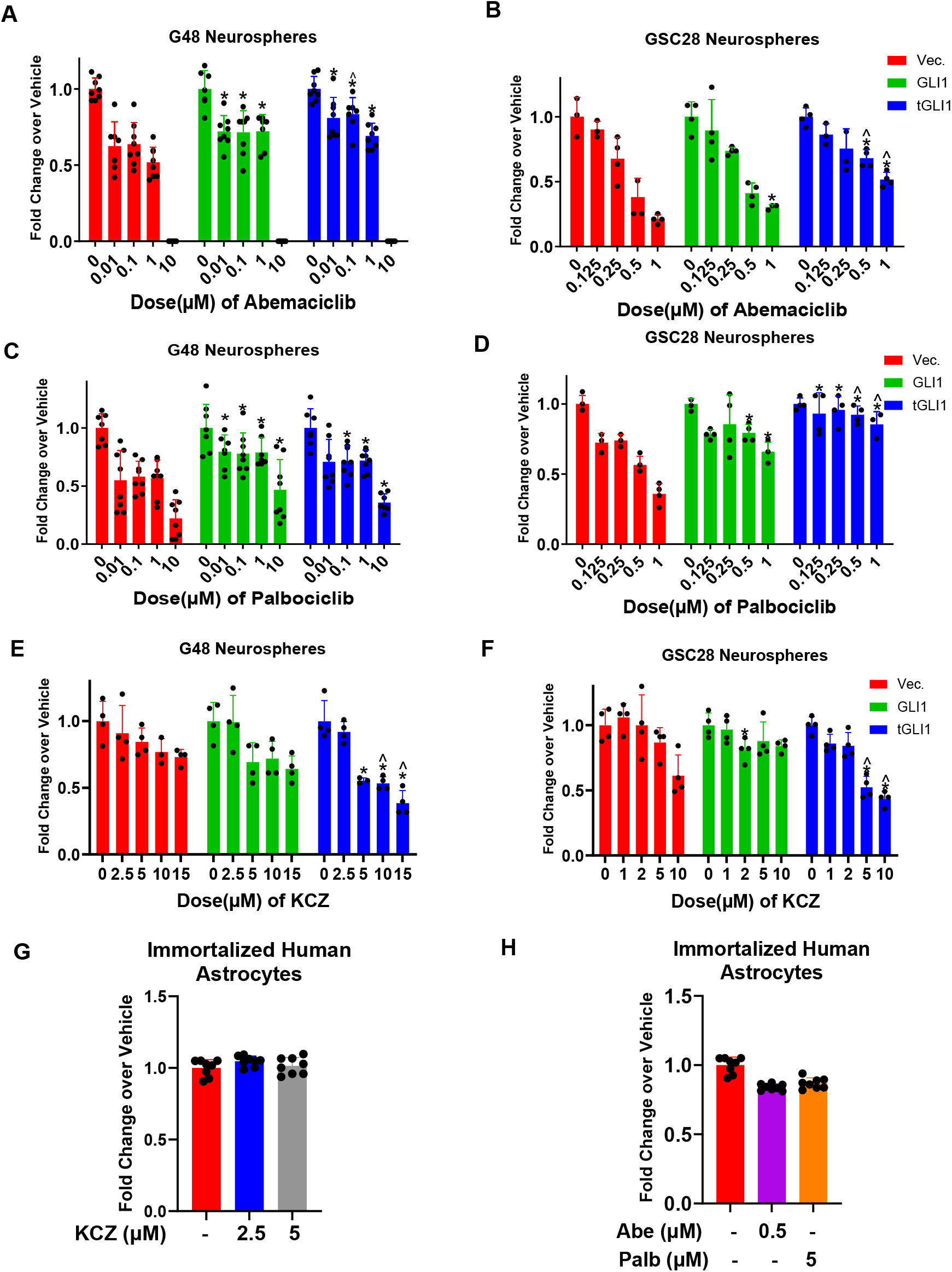
tGLI1 promotes GBM resistance to CDK4/6 inhibitors *in vitro*. **(A-B)** Isogenic G48 (A) or GSC-28 (B) vector, GLI1, and tGLI1 neurospheres were treated with increasing doses of abemaciclib. tGLI1 lines showed increased resistance to abemaciclib treatment. **(C-D)** Isogenic G48 (C) or GSC-28 (D) vector, GLI1, and tGLI1 neurospheres were treated with increasing doses of palbociclib. tGLI1 lines showed increased resistance to palbociclib treatment. **(E-F)** Isogenic G48 (E) or GSC-28 (F) -vector, -GLI1, and -tGLI1 neurospheres were treated with increasing doses of KCZ. KCZ selectively inhibited tGLI1 neurosphere formation. **(G-H)** Immortalized human astrocytes were treated with KCZ (2.5 and 5 µM) (G), Palbociclib (0.5 µM) or abemaciclib (5 µM). Treatment of all three drugs showed no signs of decreased cell viability (H). Student’s t-test was used to compute p-values. *p<0.05 compared to vector group at the same does; ^p<0.05 compared to GLI1 group at the same dose.

KCZ, an FDA approved anti-fungal, has previously been shown to specifically inhibit tGLI1 in breast cancer brain metastasis (29). Since KCZ permeates the BBB (29), we next investigated the efficacy of KCZ in tGLI1-overexpressing GBM lines. We observed that KCZ selectively inhibited GSC formation in both G48-tGLI1 and GSC-28-tGLI1 lines as compared to vector and GLI1 controls (**Fig. 3E-F**). Immortalized human astrocytes were treated with KCZ (**Fig. 3G**) or CDK4/6is (**Fig. 3H**) to determine potential treatment toxicity to brain parenchymal cells and no significant effects were observed on cell viability. Together, these results suggest that tGLI1 promotes CDK4/6 inhibitor resistance, and KCZ effectively inhibits tGLI1.

#### KCZ+Abemaciclib combination therapy significantly reduces GSCs and GBM stemness in vitro

While monotherapies have shown promise in various cancer types, combination therapies have exhibited greater effectiveness. In our study, we have shown the co-upregulation of the tGLI1 and CDK4/6 pathway in GBM, suggesting a potential therapeutic benefit through co-inhibition (**Fig. 2**). Thus, we next investigated the novel combination of KCZ and CDK4/6is, abemaciclib or palbociclib. G48-tGLI1 and GSC-28-tGLI1 cells were grown as neurospheres and were treated with vehicle, KCZ, abemaciclib/palbociclib, or combination at varying therapeutic ratios. Combination indices (CI) at different effective doses (EDs) were generated by the Chou and Talalay method, with CI<1 indicating synergism, while CI>1 indicates antagonism (**Fig. 4A, B**) (37). Combination treatment of KCZ and abemaciclib at a 5:1 ratio was synergistic in G48-tGLI1 neurospheres and significantly inhibited sphere formation compared to vehicle and single treatment (**Fig. 4A, C**). Combination ratios of 10:1, 5:1, and 2:1 were synergistic against GSC-28 tGLI1 spheres, significantly reducing sphere formation compared to vehicle and single treatment groups (**Fig. 4A, E**). Similar results were found when combining KCZ with Palbociclib (**Fig. 4B, D, F**). These results demonstrate that KCZ-mediated inhibition of tGLI1 could potentially sensitize GBM to CDK4/6i treatment.

**Figure 4:**
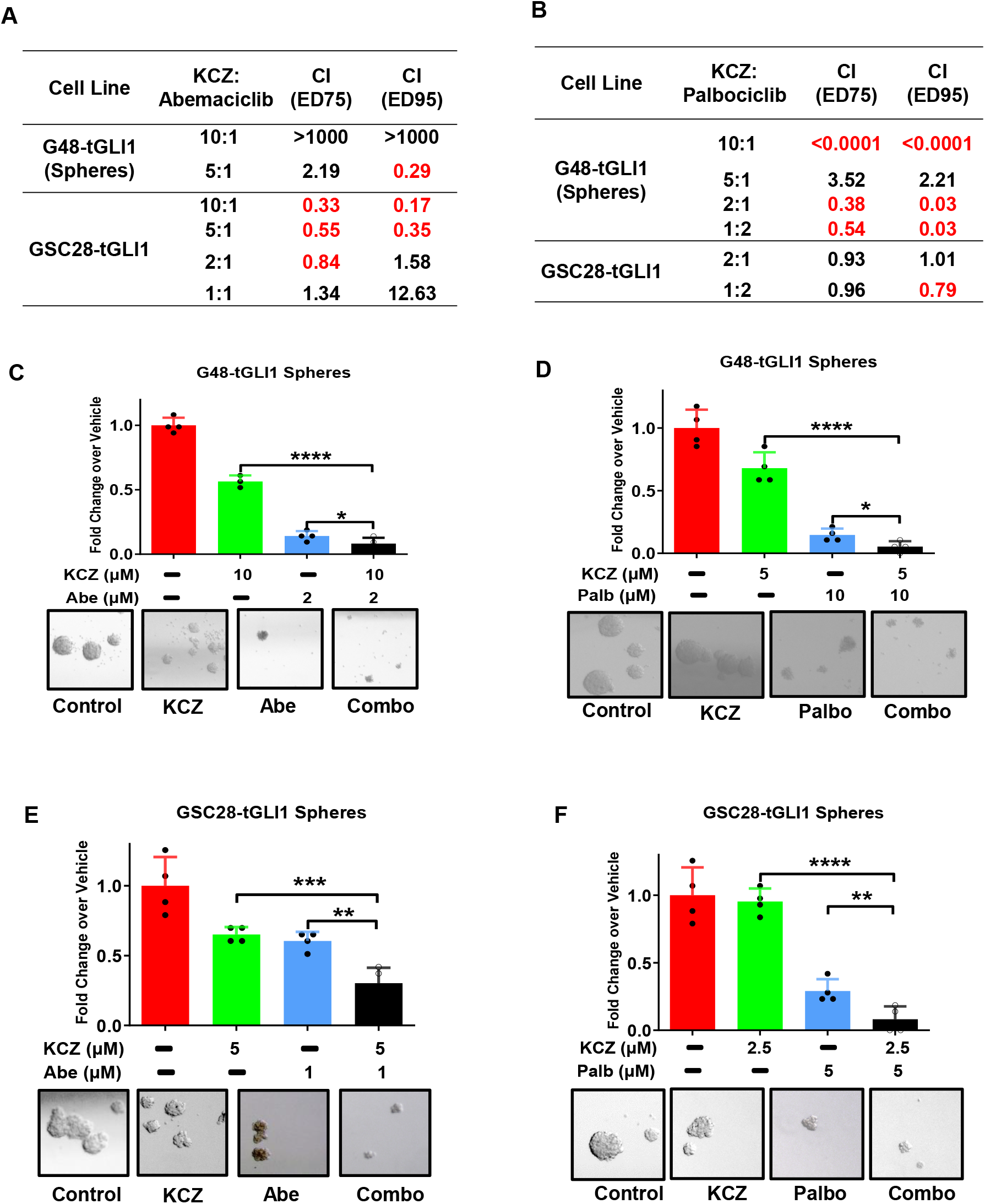
KCZ+abemaciclib Combination therapy significantly reduces GSCs and GBM stemness *in vitro*. **(A-B)** G48-tGLI1 and GSC-28-tGLI1 lines were grown as neurospheres and treated with combination therapy of KCZ+abemaciclib (A) or KCZ+palbociclib (B) at multiple combination ratios, as depicted in the respective tables. Combination index (CI) was computed using Calcusyn based on the results of neurosphere formation assay. CI<1 means synergism, CI=1 means addition, and CI>1 means antagonism. Synergistic combination ratios are highlighted in red. **(C-F)** Representative neurosphere results of synergistic combination ratios in both G48-tGLI1 and GSC-28-tGLI1 lines. Student’s t-test was used to compute p-values. Representative comparisons are labelled *p<0.05, **p<0.01, ***p<0.001, or ****p<0.0001 between two groups.

### KCZ+Abemaciclib combination therapy significantly decreases GBM proliferation and promotes apoptosis in vitro

Treatment with CDK4/6is results in cell cycle arrest in breast cancer [38]. Therefore, we next evaluated the efficacy of single or combination treatment in inducing cell cycle arrest in G48-tGLI1 and GSC28-tGLI1 lines by staining with 7-AAD, a fluorescent chemical compound to label DNA (38) (**Fig. 5A-B, Supp. Figs 7 and 8**). Results showed that KCZ treatment alone had no significant effect on altering cell cycle progression in G48-tGLI1 cells (**Fig. 5A**). However, treatment with abemaciclib and the combination significantly promoted G1 arrest (**Fig. 5A**). Interestingly, only combination treatment significantly promoted cell cycle arrest in G2 in the GSC-28-tGLI1, demonstrating the effectiveness of the combination therapy (**Fig. 5B**). The difference in treatment response across the two cell lines may be due to their different Rb status. The GSC-28-tGLI1 cells are Rb deficient and thus lack regulation of E2F1 and FOXM1 that subsequently contributes to CDK4/6i resistance, which is overcome with combination therapy with CDK4/6i and KCZ.

**Figure 5:**
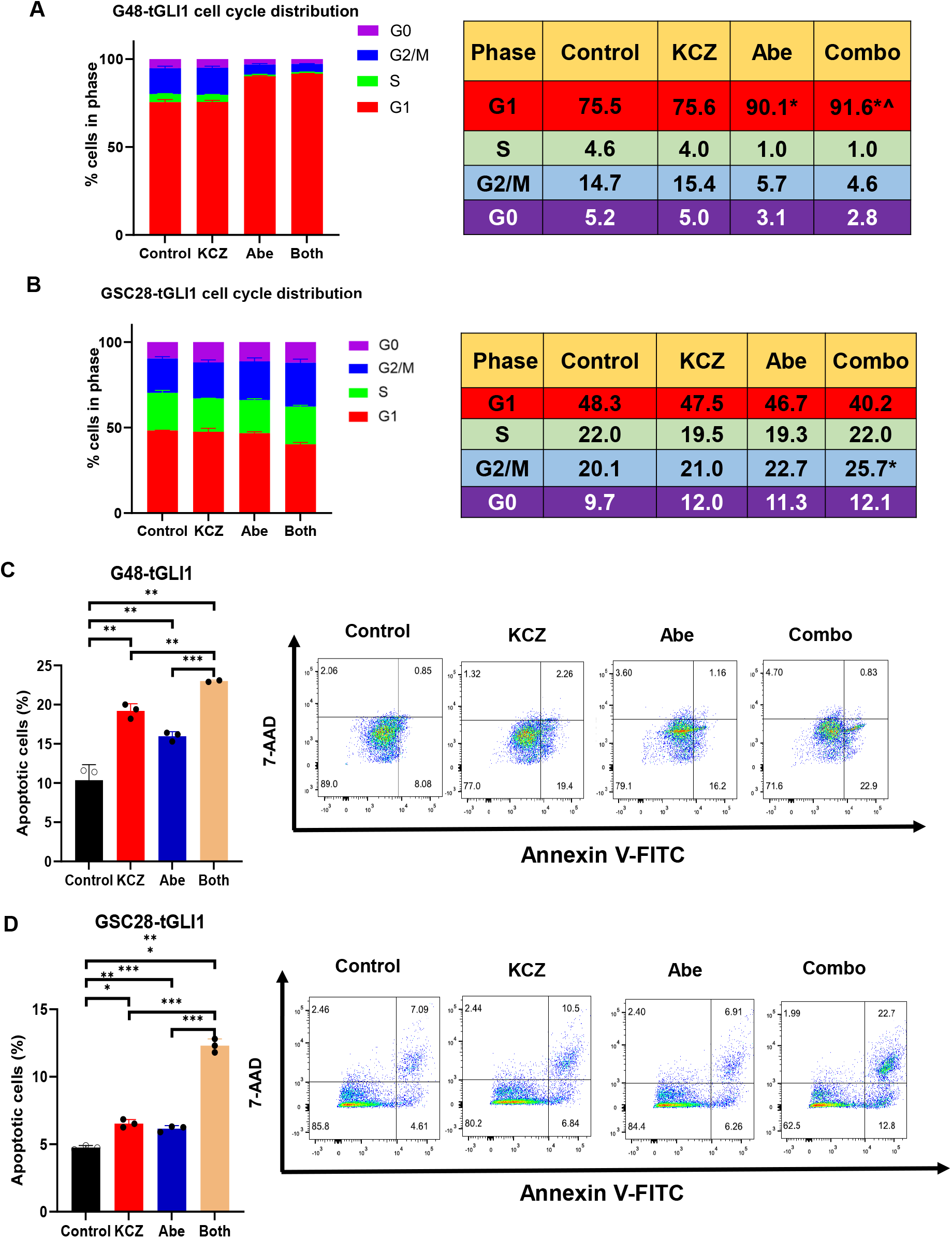
KCZ+abemaciclib combination therapy significantly decreases GBM proliferation and promotes apoptosis *in vitro*. **(A-B)** Combination therapy significantly inhibits cell cycle progression in G48-tGLI1 (A) and GSC-28-tGLI1 (B) cell lines. Cells were treated with vehicle, KCZ (5 µM), abemaciclib (1µM), or KCZ+abemaciclib combination for 24 hours. Student’s t-test was used to compute p-values. *p<0.05 compared to control in %cells of G1 phase, ^p<0.05 compared to abemaciclib-only group in % cells of G1 phase. **(C-D**) G48-tGLI1 (C) or GSC-28-tGLI1 (D) cells treated with vehicle, KCZ (5µM), abemaciclib (1µM), or KCZ+abemaciclib combination for 24 hours and then subjected to Annexin V-FITC/7-AAD flow cytometry. Combination therapy significantly promoted apoptosis in both cell lines. Representative flow cytometry diagrams are shown on the right. Student’s t-test was used to compute p-values. *p<0.05, **p<0.01, ***p<0.001 between two groups.

Cell cycle arrest often leads to cell death, therefore, we next investigated changes in apoptosis following single or combination therapies (39). G48-tGLI1 and GSC-28-tGLI1 cell lines were subjected to vehicle, single agent, or combination therapy for 24 hours, and then stained with Annexin V-FITC/7-AAD and analyzed by flow cytometry analysis to assess apoptosis (**Fig. 5C-D**). Results showed that while both single treatments significantly increased apoptosis in G48-tGLI1 cells as compared to vehicle, combination treatment significantly increased apoptosis compared to the other two groups(**Fig. 5C**). Similar results were observed in GSC-28-tGLI1 (**Fig. 5D**). These results further confirm the synergy between tGLI1 and CDK4/6 inhibition in GBM, by promoting cell cycle arrest and increasing GBM apoptosis *in vitro*.

### KCZ+Abemaciclib Combination therapy significantly decreases tumor growth in vivo

To assess the translatability of our findings, we investigated the efficacy of this combination therapy *in vivo*. G48-tGLI1 cells containing a luciferase reporter were intracranially implanted into the right frontal lobe of female athymic nude mice. After a 10 day period of tumor engraftment, animals were randomized, after which mice were treated with vehicle, KCZ (50 mg/kg, intra-peritoneal) only, abemaciclib (50 mg/kg, oral gavage) only, or KCZ+abemaciclib combination therapy for ∼28 days (n=9-10 mice/ group). Combination therapy significantly inhibited tumor growth as compared to control, while single treatment had no significant effect (**Fig. 6A-B, Supp Fig.9**). All treatment groups were well tolerated as determined through an alanine transaminase assay (ALT) to determine liver toxicity and monitoring of mouse weight over the course of the study (**Fig. 6C-D**).

**Figure 6:**
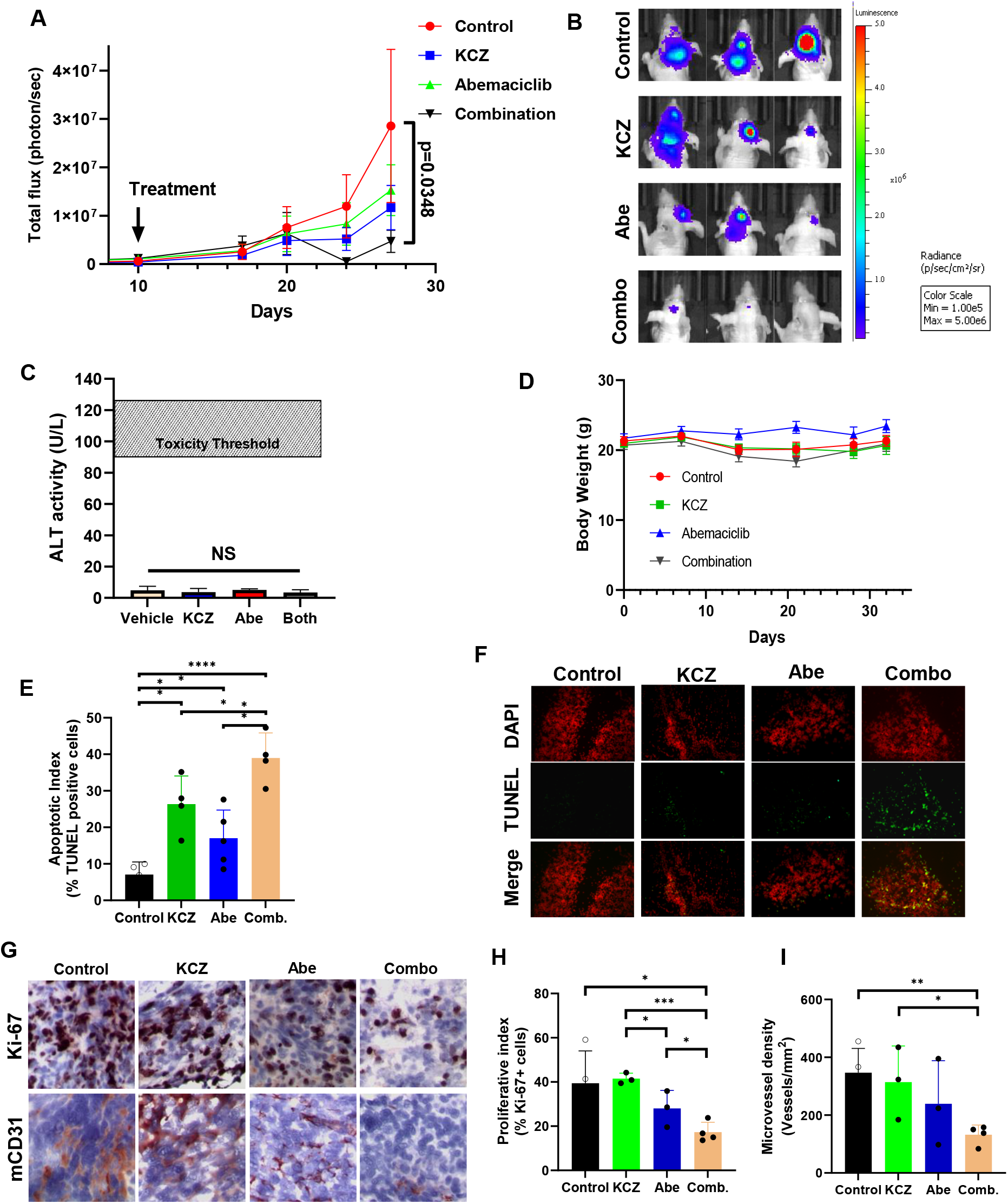
KCZ+abemaciclib combination therapy significantly decreases tumor growth *in vivo*. **(A)** Combination treatment significantly reduced tumor growth in mice. Growth curve of G48-tGLI1 intracranial tumors across control, KCZ, abemaciclib, and combination groups. **(B)** Representative IVIS images at 27 days after intracranial inoculation. **(C)** Blood ALT levels were not significantly increased in any treatment group. **(D)** No significant change in mouse weight was seen across all treatment groups. **(E)** Combination treatment significantly increased GBM apoptosis *in vivo*. TUNEL staining conducted on sectioned mouse brains across control, KCZ, abemaciclib, and combination groups. **(F)** Representative TUNEL staining images. **(G)** IHC staining of sectioned mouse brains stained for Ki-67 and mCD31. Combination treatment significantly decreased GBM proliferation and microvessel formation *in vivo*. **(H-I)** Corresponding graphs for Ki-67 (H) and mCD31 (I) IHC staining. Student’s t-test was used to calculate p-values.

Mouse brains were harvested to be analyzed for changes in tumor proliferation, angiogenesis, and cell death following treatment using IHC. The combination therapy significantly increased Terminal deoxynucleotidyl transferase dUTP nick-end labeling (TUNEL) positive cells as compared to vehicle and single treatment groups, demonstrating increased tumor apoptosis (**Fig. 6E-F**). Furthermore, treatment with both agents significantly increased cell death as compared to vehicle (**Fig. 6E-F**). Additionally, mouse brain sections were stained with Ki-67 and mCD31 to determine changes in tumor proliferation and microvessel formation, respectively. Results showed a significant decrease in Ki-67 positivity following combination therapy compared to vehicle and single treatment groups, while either single treatment had no significant change compared to vehicle (**Fig. 6G-H**). Furthermore, microvessel formation was significantly inhibited by combination therapy when compared to vehicle and KCZ alone groups (**Fig. 6G and I**). These results demonstrate that the KCZ+abemaciclib combination therapy retains their synergistic effect *in vivo*, significantly inhibiting GBM growth and promoting apoptosis.

### KCZ+Abemaciclib combination therapy inhibits PDX39 growth in an intracranial mouse model

To further validate the efficacy of KCZ+abemaciclib combination therapy against GBM in a more clinically relevant *in vivo* model, we utilized a second orthotopic xenograft mouse model using the highly aggressive PDX 39 line that is positive for tGLI1 and CDK4/6 expression (**Fig. 2J**). Female athymic nude mice were intracranially inoculated with PDX39 cells containing a luciferase reporter into the right frontal lobe and animals were treated with vehicle, KCZ (50 mg/kg), abemaciclib (50 mg/kg), or combination therapy for 20 days (**Fig. 7A**). Following termination, brains were harvested and underwent *ex vivo* imaging, where results showed a significant reduction in brain bioluminescence in the combination therapy group, as compared to all other treatment groups (**Fig. 7B-C**). Additionally, there was no significant change in mouse weight over the course of treatment, indicating that treatment was well tolerated (**Fig. 7D**). These results, along with the results from our previous animal study (**Fig. 6**), highlight the efficacy and safety of the synergistic KCZ+abemaciclib combination therapy *in vivo*.

**Figure 7:**
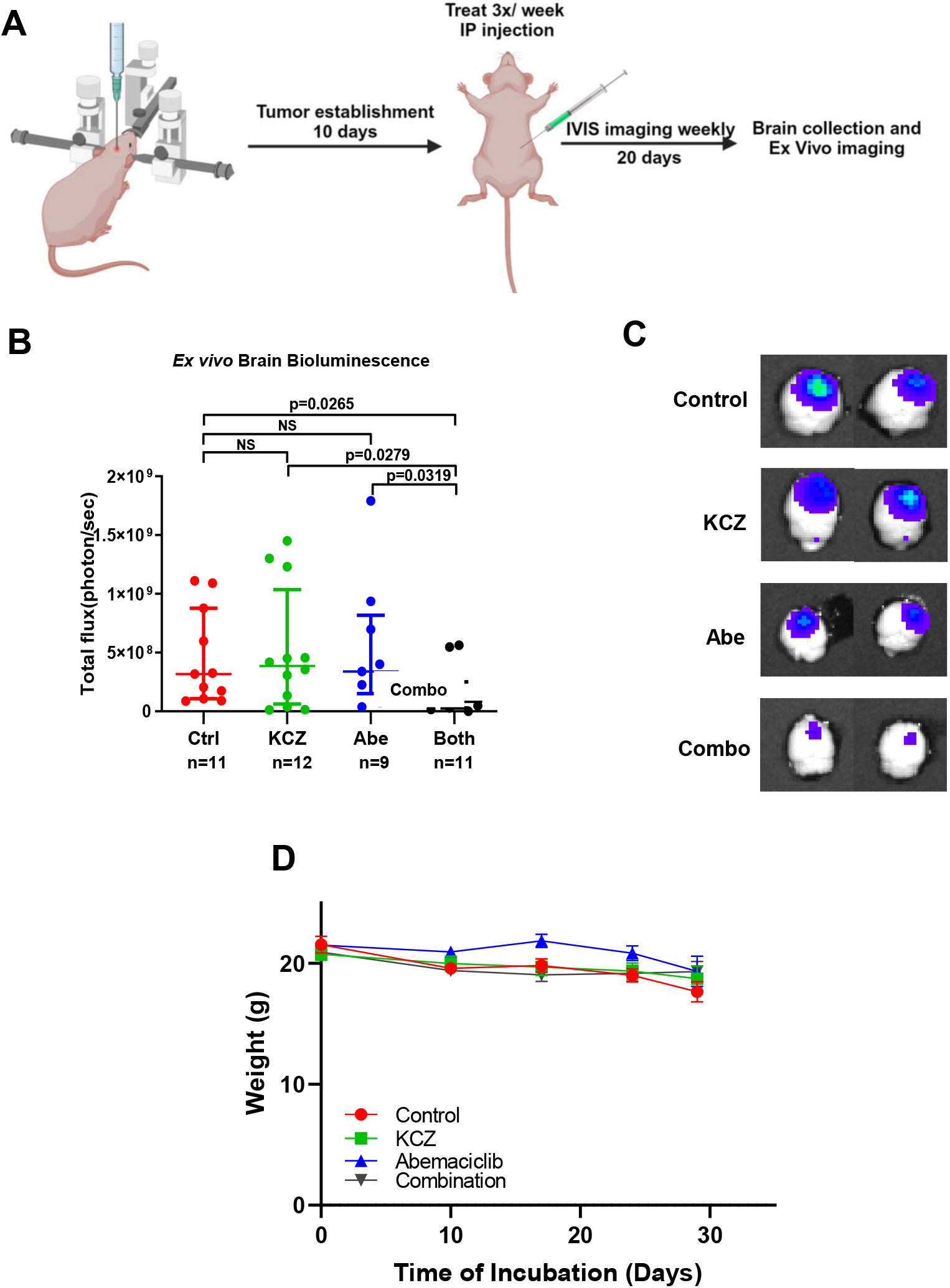
KCZ+abemaciclib combination therapy inhibits PDX39 growth in an intracranial mouse model. **(A)** Animal study schematic. Mice were intracranially inoculated with PDX39 luciferase expressing cells in the right frontal lobe. Tumors were allowed to establish for 10 days before randomization and the beginning of treatment. The study was terminated 30 days after inoculation. **(B)** *Ex vivo* brain bioluminescence of control, KCZ, abemaciclib, and combination groups. Combination therapy significantly reduced PDX39 growth *in vivo*. **(C)** Representative *ex vivo* brain images. **(D)** No significant change in mouse weight was seen across all treatment groups. Student’s t-test was used to calculate p-values.

#### tGLI1 promotes resistance to GBM standard-of-care therapy and KCZ overcomes this resistance

Radiation therapy is a major component of the standard-of-care therapy for GBM patient, however, GSCs have been shown to promote radiation resistance (30, 40, 41). We have previously shown tGLI1 promotes the GSC population, therefore, we next investigated tGLI1 expression and GBM radiation resistance (26-28). Isogenic G48-vector, -GLI1, and -tGLI1 stable lines were exposed to 0 gray (Gy) or 5 Gy of ionizing radiation (IR), and were then seeded in monolayer or sphere forming conditions (**Fig. 8A-B**). Interestingly, tGLI1 overexpression had no effect on G48 cells grown in monolayer conditions, but significantly promoted survival in neurosphere following IR **(Fig. 8A-B; Supp. Fig. 10**). Similar results were seen with GLI1 and tGLI1 overexpressing GSC-28 neurospheres, which are more resistant to 2 Gy of IR treatment as compared to GSC-28-vector cells **(Fig. 8C)**.

**Figure 8:**
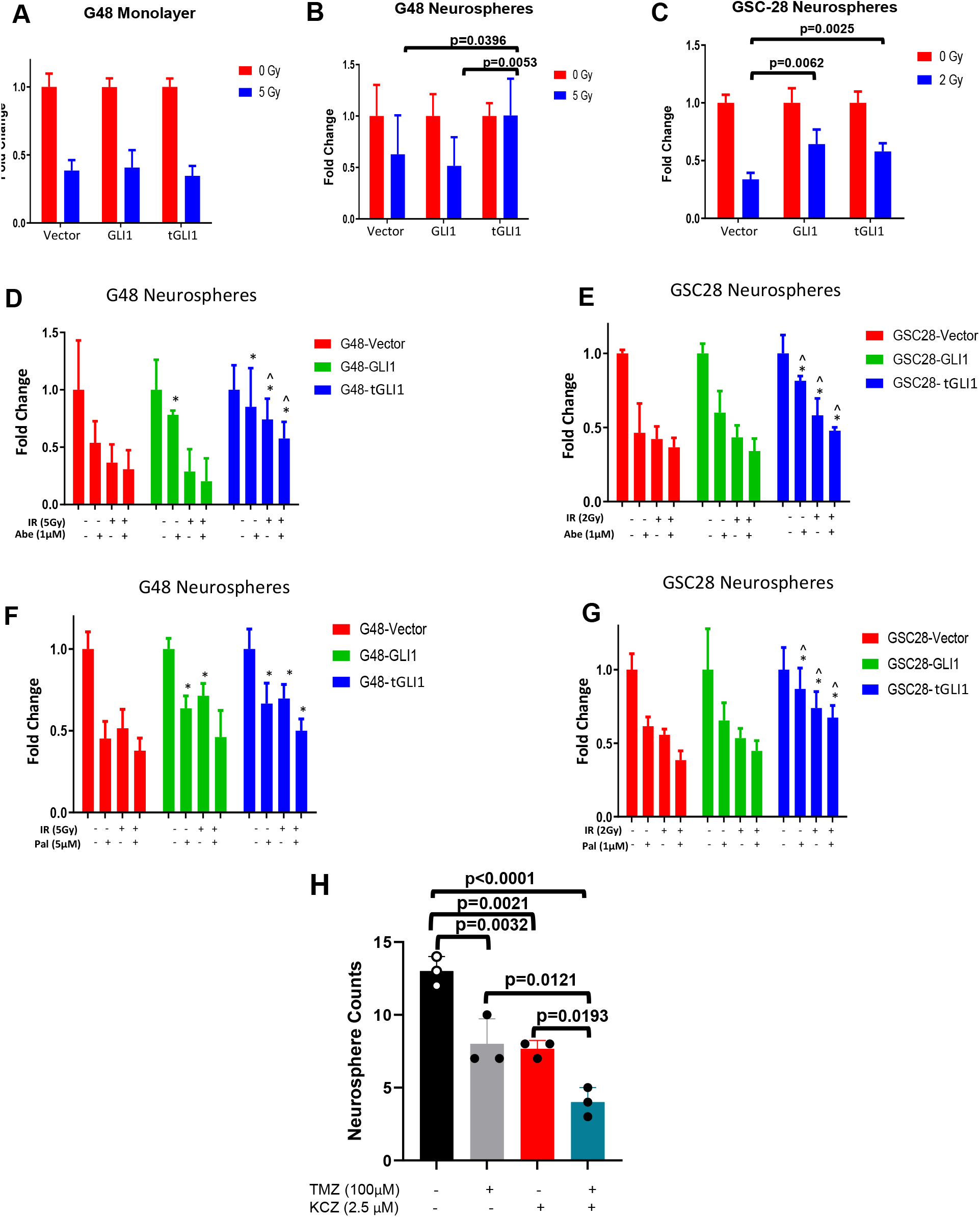
tGLI1 promotes resistance to GBM standard-of-care therapy and KCZ overcomes the resistance. **(A-B)** tGLI1 overexpression promoted radioresistance in isogenic G48 neurospheres, but not monolayer. Isogenic G48 vector, GLI1, and tGLI1 were treated with 0 Gy or 5 Gy of radiation therapy (RT), and then seeded in monolayer (A) or sphere forming conditions (B). **(C)** GSC-28-GLI1 and tGLI1 lines were more resistant to RT. Isogenic GSC-28 vector, GLI1, and tGLI1 cells were treated with 0 Gy or 2 Gy of RT, and then seeded in sphere forming conditions. **(D-E)** G48-tGLI1 and GSC-28-tGLI1 lines are resistant to RT, abemaciclib, and RT+abemaciclib combination treatment compared to vector and GLI1 lines. Isogenic G48 D) or GSC-28 (E) vector, GLI1 and tGLI1 cells were treated with vehicle, RT (5 Gy forG48; 2 Gy for GSC-28), abemaciclib (1 µM), or RT+abemaciclib combination and seeded in sphere forming conditions. **(F-G)** G48-tGLI1 and GSC-28-tGLI1 lines were resistant to RT, Palbociclib, and RT+palbociclib compared to vector and GLI1 lines. Isogenic G48 (D) or GSC-28 (E) -vector, -GLI1 and -tGLI1 cells were treated with vehicle, RT (5 Gy for G48; 2 Gy for GSC-28), Palbociclib (5 µM for G48; 1 µM for GSC-28), or RT+palbociclib combination and seeded in neurosphere forming conditions. **(H)** G48-tGLI1 cells had increased sensitivity to TMZ treatment when combined with KCZ treatment. G48-tGLI1 cells were grown in sphere forming conditions and treated with vehicle, TMZ (100 µM), KCZ (2.5 µM), or TMZ+KCZ combination therapy and spheres were counted. Student’s t-test and One-way ANOVA was used to calculate p-values.

Next, we investigated changes in neurosphere forming abilities following radiation alone (5 Gy for G48, 2 Gy for GSC-28), CDK4/6i treatment alone, or combination in G48 and GSC-28 isogenic stable lines. Interestingly, we found tGLI1-overexpressing G48-tGLI1 and GSC-28-tGLI1 cells to be more resistant to radiation alone, abemaciclib (1µM in both cell lines) alone, and combination therapy as compared to their respective vector and GLI1 lines (**Fig. 8D-E**). Similar results were seen with the use of Palbociclib (5µM for G48 and 1µM for GSC-28) in G48 and GSC-28 isogenic stable lines (**Fig. 8F-G**). Together, these results demonstrate that tGLI1 promotes radiation resistance, specifically in the GSC population. This is further supported by the observation that combined radiation and CDK4/6i treatment is less effective against tGLI1-overexpressing GBM in inhibiting neurosphere formation.

In addition to radiation, TMZ treatment is a second component of the standard-of-care therapy for GBM patients (3). We further investigated the role of tGLI1 in TMZ resistance in GBM. We treated isogenic G48-tGLI1 spheres with vehicle, 2.5 µM KCZ alone, 100 µM TMZ alone, or KCZ+TMZ combination therapy and counted neurospheres on day 5 following the initial treatment. Results showed KCZ and TMZ monotherapy significantly reduced sphere formation compared to vehicle (**Fig. 8H**). Importantly, the combination treatment was the most effective in reducing neurosphere formation compared to all other groups (**Fig. 8H**). Together, results in Fig. 8 demonstrated that tGLI1 promotes resistance to GBM standard-of-care therapy and KCZ overcomes the resistance

## DISCUSSION

CDK4/6is have shown promise in effective treatment of different cancer types and are now in clinic with FDA approval for treating hormone receptor-positive breast cancers and tumors with loss of p16^INK4a^ or increased cyclin D1 (42). Unfortunately, clinical trials in GBM patients have not yielded positive results in improvement of OS or progression-free survival (PFS) owing to therapeutic resistance in this tumor type (13, 21, 22). Following investigation of underlying mechanisms of resistance, we report that tGLI1 promotes GBM resistance to CDK4/6i. We also found that tGLI1 overexpression promotes tumor resistance to RT and TMZ that are postoperative standard-of-care therapies for GBM. Recent work in our lab has demonstrated that tGLI1 is enriched in mGBM, an aggressive subtype of GBM that exhibits characteristics of increased stemness, drug resistance, and tumor invasion (26, 35, 43). Moreover, our group has also shown that tGLI1 plays a critical role in enriching the GSC population by promoting mGBM. Other studies have shown that the GSC population plays a pivotal role in treatment resistance to RT, chemotherapies, and other targeted therapies such as bevacizumab in GBM (30, 35). Thus, we speculate that one of the ways tGLI1 leads to treatment resistance is by increasing glioma stemness (23, 27). However, there may be multiple pathways interplaying in the tGLI1-mediated resistance to CDK4/6is in GBM. Previous work from our group reported that tGLI1 overexpressing GBM cells have increased murine double minute 2 (MDM2) expression, a reported factor that promotes GBM CDK4/6i resistance (23, 44). Moreover, we have previously identified the novel interaction between tGLI1 and STAT3, with STAT3 also being reported to promote CDK4/6i resistance in breast cancer (27, 45). However, the role of these two factors in mediating CDK4/6i resistance in the context of tGLI1-overexpressing GBM has not been studied and warrants further investigations.

Another key aspect of our work is the demonstration of increased treatment efficacy with a combination therapy of CDK4/6is and KCZ, which strongly suggests the therapeutic value of targeting tGLI1 in GBM to unleash the full potential of CDK4/6is. tGLI1 is a terminal effector of the SHH pathway, one of the pathways that significantly correlated with CIRS. Two additional pathways, namely, β-catenin and PI3K pathways, are also significantly correlated with CIRS. In a recent study, combination of Wnt/β-catenin inhibitor (AC1Q3QWB) with palbociclib was synergistic against GBM in mouse models (46). Additionally, combination of PI3K/AKT/mTOR pathway inhibitors sensitized breast cancer lines to CDK4/6is, with similar results being seen in GBM (47, 48). Furthermore, combining CDK4/6is with immunotherapies have shown increased efficacy in breast cancer models and brain metastasis studies (49, 50). Taking these studies into consideration, there is a need for further investigation of novel combination targeted therapeutics that can overcome treatment resistance in GBM.

Our findings have significant translational implications in overcoming tumor treatment resistance to CDK4/6is, RT, and TMZ and preventing tumor progression while effectively treating GBM using a novel combination therapy. In patient specimens and *in vitro* GBM models, we found that tGLI1 pathway activation is highly correlated with the CDK4/6 pathway. Moreover, we demonstrated that tGLI1 promotes CDK4/6i resistance in GBM and that dual inhibition of tGLI1 and CDK4/6 with KCZ and CDK4/6is has synergistic treatment effects *in vitro* and *in vivo*. Also, tGLI1 promotes GBM resistance to RT, with modest effects in sensitization to RT when treated with CDK4/6 inhibitors. Lastly, we showed that KCZ sensitizes GBM to TMZ treatment prolonging the treatment window with standard-of-care chemotherapy. All-in-all, our study demonstrates that tGLI1 is a novel targetable transcription factor contributing to CDK4/6i and chemoradiation resistance in GBM. Importantly, congruent to our findings in metastatic breast cancer to the brain, we also showed that KCZ, FDA-approved antifungal agent that crosses the BBB, is a viable therapeutic agent that selectively binds to tGLI1 and overcomes CDK4/6i resistance in GBM (29). Thus, treating GBM with a combination therapy of KCZ and CDK4/6is may maximize the full potential of CDK4/6i in treating GBM. Further investigation is warranted in clinical trials to evaluate the efficacy of this drug combination in the clinic.

## METHODS

### Establishment of Stable Isogenic Cell Lines

The isogenic G48LL2 and GSC28 cell lines stably expressing vector, GLI1 or tGLI1, were generated as previously described (27).

### Cell Viability Assay

Cells were seeded at a concentration of 2,000 cells/well in 96-well white bottom plates (Greiner Bio-One; Kremsmünster, Austria), and incubated at 37°C, 5% CO^2^ for 24 hours. Cells were treated with test compounds for 48 hours. Viability was determined using the CellTiter-Blue Cell Viability Assay (Promega; Madison, WI, USA) according to the manufacturer’s instructions. Fold changes over vehicle at various doses were computed.

### Western Blot

Western blots were performed as previously described (6). Antibodies used for western blot included CDK4 (Cell Signaling Technology (CST); #12790), CDK6 (CST; #13331), Rb (Cell signaling; 9309), E2F1 (CST; #3742), Cyclin D1 (CST; 2922), FOXM1 (CST; #20495), α-tubulin (Sigma-Aldrich; T6074), βactin (Sigma-Aldrich; T4026), GLI1 (CST; #2643), and the rabbit polyclonal tGLI1 antibody that was developed and validated by our laboratory (25, 28, 29, 43).

### TUNEL Assay

TUNEL assay was used to determine the extent of apoptosis. Xenograft-bearing brain samples were embedded in OCT medium and were sectioned using a cryostat. Brain sections were subjected to TUNEL assay using the Click-iT Plus TUNEL Assay (Invitrogen) according to the manufacturer’s instructions. Following nuclear counterstaining by DAPI (Invitrogen), the slides were imaged by fluorescence microscope (Zeiss). Apoptotic index was generated by calculating the percentage of TUNEL positive cells for each animal.

### Flow Cytometry Analysis

For apoptosis analysis, cells were harvested and washed with PBS after 24-hour treatment. A total of 1.5×10^6^ cells per group were suspended in 1mL 1X binding buffer from Annexin V-FITC Apoptosis Staining / Detection Kit (abcam, #14085), after which cells were stained with 10μL Annexin V-FITC and 10μL 7-AAD (abcam, #228563, 50 μg/mL). For cell cycle phase distribution analysis, the cells were harvested and washed by PBS after 24-hour treatment. A total of 6×10^6^ cells per group were fixed by ice-cold 70% ethanol. The cells were stained by 7-AAD (25 μg/mL) after two PBS washes. Stained cells for apoptotic or cell cycle analysis were detected by BD LSRFortessa X-20 Analyzer within 2 hours. The data were collected and analyzed by FlowJo™ software (BD Biosciences).

### In vivo studies

Female athymic nude (nu/nu) mice (5-6 weeks old; Charles River Laboratories; Wilmington, MA) were used. All mice were housed in a pathogen-free facility of the Animal Research Program at Wake Forest School of Medicine (WFSM) under a 12:12-hour light/dark cycle and were fed with sterile rodent chow *ad libitum*. Animal handling and procedures were approved by the WFSM Institutional Animal Care and Use Committee (IACUC) (protocol #A20-076). A total of 1 × 10^5^ G48 cells stably expressing a tGLI1 expression vector (G48-TGLI1) in 5 μL PBS were injected into the right frontal lobe of anesthetized female athymic nude (nu/nu) mice (5-6 weeks old). Cells were allowed to engraft for 10 days. Then, mice were randomized into control (100% PEG-300), 50 mg/kg KCZ, 50 mg/kg abemaciclib, or the combination therapy groups according to the visualization of brain bioluminescent signal. KCZ was formulated in 100% PEG-300 and was administered three times per week by intraperitoneal (IP) injection. abemaciclib was formulated in 100% PBS and was administered every day by oral gavage. Tumor progression was monitored by biweekly bioluminescent imaging during which xenograft bearing mice were injected IP with 100 mg/kg D-luciferin (Perkin Elmer) and imaged using the IVIS Lumina LT Series III imager (Perkin Elmer) after 5-minute incubation. Tumor burden was analyzed by quantifying total bioluminescent signal in brain by the Living Image software version 4.7.2 (Perkin Elmer).

### Statistical Analysis

The values were presented as mean ± SD unless noted otherwise. Student’s t-test, Log-rank test, Two-way ANOVA and Fisher’s exact test were performed to determine the difference between groups using GraphPad Prism 9 or Microsoft Excel.

## Supporting information

Original Western Blots

## Authors Contributions

Conceptualization: Y.Y. and H.W.L. Original draft writing, preparation of figures and tables: Y.Y., A.A., A.C. (co-first authorship). Data acquisition and analysis: Y.Y., A.A., A.C., M.C., D.Z., A.D.O. Revising and editing: A.C., C.Z., M.K.N., M.S.K., D.Z, A.D.O., Y.E., N.T., J.J.Z., S.H.H., R.E.S., M.C., H.W.L. Supervision; H.W.L. All authors have read and agreed to the published version of the manuscript.

## Acknowledgements

We acknowledge funding support for this study from NIH grants R01CA228137 (H-WL), R21CA286225 (H-WL) and UE5NS113757 (AC and NT)(Research Education Programs for Residents & Fellows), DoD grants W81XWH-19-1-0072 (H-WL), W81XWH-20-1-0044 (H-WL), W81XWH-19-1-0753 (H-WL), and HT9425-24-1-0889 (H-WL), as well as, MetaVivor Translational Research Grant (H-WL) and UTStars (H-WL).

