## Supplementary Methods for "Targeting tGLI1, a novel mediator of tumor therapeutic resistance, using Ketoconazole sensitizes glioblastoma to CDK4/6 therapy and chemoradiation"

### *Reverse Transcription-Quantitative Polymerase Chain Reaction (RT-qPCR)*

The isogenic G48 and GSC28 cell lines stably expressing control vector, GLI1 and TGLI1 were harvested and total RNA was extracted using RNeasy Plus Mini Kit (Qiagen) according to the manufacturer's instructions. Complementary DNA (cDNA) was then synthesized using the Superscript III First-Strand cDNA synthesis system (Invitrogen, #18080051). The forward and reverse primers used for regular PCR were: 5'-CTTCTGCAGTCCACATATGCAACA-3' and 5'-CAACTGGTCGGCTTCAGAGTTTC-3' (CDK4) (1), 5'-AAGTTCCAGAGCCTGGAGTG-3' and 5'-CGATGCACTACTC GGTGTGA-3' (CDK6) (2), 5'- CACCCCAGTGCCAACCGCTACTTG-3' and 5'-AAAGAGGAGCTATCCCCTCCTCAG-3' (FOXM1) (3) 5'-CACCAAGCTAACCTCAT GTC-3' and 5'-CGGGGAGAAGAAAAGAGTGGG-3' (GLI1) (4), 5'-GTGTGGGGA CAGAAGTCAA-3' and 5'-GTGCGGATAACCGTCTGC-3' (TGLI1) (4), and 5'-ACTGCCAACGTGTCAGTGG-3' and 5'-GTGTCGCTGTTGAAGTCAGA-3' (GAPDH) (4). QPCR was performed using GoTaq qPCR Master Mix (Promega). GAPDH was used as a control for gene expression normalization and three independent technical replicates were performed for each sample.

### *Extraction and Cryopreservation of Brains*

Mice were euthanized by IP injection with 10 mL/kg of a mixture of 130 mg/kg ketamine and 10 mg/kg xylazine. Chemical euthanasia was confirmed by cervical dislocation. Following decapitation, mice brains with tumor xenografts were rapidly excised. Following PBS washes, brains were embedded in NEG-50 Frozen Section Medium (OCT medium, Richard-Allen Scientific, 6502) and stored at -80°C until sectioning.

#### *Alanine Transaminase (ALT) Assay*

ALT assay was performed to determine alanine transaminase activity in mouse serum using the Alanine Transaminase Colorimetric Assay kit (Cayman Chemical) to monitor acute liver toxicity. Following sacrifice, mice were immediately exsanguinated by intracardiac puncture. Serum was separated by placing blood samples at room temperature for 30 minutes followed by a centrifugation at 2000 x g for 15 minutes. Serum samples were then subjected to ALT assay according to the manufacturer's instructions. After setting up the plate including a positive control, absorbance at 340 nm was measured every minute for 10 minutes at 37°C using a SpectraMax iD3 (Molecular Devices; San Jose, CA, USA). The change in absorbance (ΔA340) per minute was computed for each 14 sample. ALT activity (U/L) was then determined based on the change in absorbance (ΔA340) per minute according to the manufacturer's instructions.

#### *Bioinformatics Analysis of GBM Database*

A gene expression profile containing 165 GBM samples with subtype and overall survival information was compiled based on The Cancer Genome Atlas (TCGA). A tGLI1 activation signature (tGAS) was established by integrating six novel tGLI1 target genes (CD24, VEGFA, VEGFC, HPA1, TEM7, VEGFR2) according to our previous research [4]. A signature including 11 genes to assess CDK4/6 inhibitor resistance (CIRS) was developed and validated by a research team in Université Libre de Bruxelles (5). Gene expression signatures of various pathway activity (β-catenin, SHH, PI3K, MTORC1, P38, MYC, PDGF, TGFB, NEDD4, AKT, WNT, RAC1, ERK1-2, JNK, MAPK, EGFR, KRAS, STAT3, NOTCH1, VEGFR1-2, TNFA, NFKB) were developed by a research team in Duke University Medical Center (6). Quantification of gene expression signature was performed by a sum of median-centered expression level of all genes. Pearson correlation was performed to determine the relationships between tGAS and CIRS. Gene set enrichment analysis (GSEA) was performed to explore the pathway enrichment in high CIRS samples. Gene Matrix file (gmx) was generated based on the signatures of pathway activity above. The Gene Cluster Text file (.gct) was generated from the TCGA dataset above. The Categorical Class file (.cls) was generated based on CIRS scores. A histogram for the score of CIRS was used to determine the threshold for high or low expression of 10 CIRS in GSEA. The cutoff was upper 33%. The number of permutations for GSEA was set to 1,000. CBioPortal online database was used to analyze CDK4/6 amplification and p15/16 or Rb deletion or mutation status in GBM samples (n=821). Kaplan-Meier curves were generated based on the patients with or without CDK4 or CDK6 amplification

***Cell lines, specimens and reagents***

Human GBM cell lines U87MG, LN18, and U251MG were obtained from ATCC and cultured according to the specified recommendations from ATCC. Luciferase expressing GBM cell line G48LL2 was developed by Dr. Waldemar Debinski (4) and were cultured in RPMI 1640 (Corning 10-041-CV) supplemented with 10% fetal bovine serum (FBS, Corning 35-10-CV) and 1% Penicillin-Streptomycin solution (P/S, Corning 30-002- CI). Patient-derived GSCs (proneural subtype: GSC-11 and GSC-23; mesenchymal subtype: GSC-20 and GSC-28) were kind gifts from Drs. Erik Sulman and Krishna Bhat at University of Texas MD Anderson Cancer Center (7). GSCs were cultured under neurosphere-forming conditions in serum-free Dulbecco's modified Eagle's medium/F12 (DMEM/F12, Gibco 11320033) supplemented with 1X B27 (Gibco 17504044), FGF

(20ng/mL, Sigma-Aldrich F0291), EGF (20ng/mL, Sigma-Aldrich E9644) and 1% Penicillin-Streptomycin solution (P/S, Corning 30-002-CI) for maintaining stem-like properties. Immortalized human astrocytes (UC1) was a kind gift from Dr. Russell O. Pieper at University of California San Francisco (8) and was cultured in Dulbecco's modification of eagle's medium (DMEM, Corning 10-013-CMR) supplemented with 10% fetal bovine serum (FBS, Corning 35-10-CV) and 1% Penicillin-Streptomycin solution (P/S, Corning 30-002-CI). Patient-derived GBM xenografts (PDX-16, PDX-39 and PDX-80) were obtained from the Mayo Clinic and maintained through serial passaging in mice flanks.

Glioma tissue microarray (GL2083) was purchased from US Biomax. CDK4/6 inhibitors abemaciclib (LY2835219, Cat. No. S5716), palbociclib (PD0332991, Cat. No. S1116) and ribociclib (LEE011, Cat. No. S7440) were purchased from Selleck Chemicals. Abemaciclib mesylate (Cat. No. 206973) for animal study was purchased from Medkoo Biosciences. Ketoconazole (KCZ, Cat. No. 15212) was purchased from Cayman Chemical.

***Immunohistochemistry (IHC)***

Tissue microarrays (GL2083) consisting of paraffin-embedded microsections were deparaffinized, dehydrated, and subjected to antigen retrieval by boiling with citrate buffer for 20 minutes. Endogenous peroxidase activity was blocked by 0.3% hydrogen peroxide. After blocking in 5% normal goat serum for 1 hour, the slides were incubated with CDK4 antibody (Cell signaling; 12790, 1:50), CDK6 antibody (abcam; 124821,1:100) or the rabbit polyclonal tGLI1 antibody (1:100) that was developed and validated by our laboratory at 4°C overnight (9-11). VECTASTAIN® ABC-HRP Kit (Vector Laboratories;

PK-4001) and ImmPACT NovaRED Substrate (Vector Laboratories; SK-4805) were used to detect primary antibodies. Following nuclear counterstaining by hematoxylin, the slides were mounted and imaged using a Pico Automatic Cell Imager (Molecular Devices). Frozen mouse brain sections (10 µm thick) were fixed by cold acetone and were subjected to endogenous peroxidase block as described above. After blocking in 5% normal goat serum for 1 hour, the slides were incubated with Ki-67 antibody (Cell signaling; 9027, 1:400) or anti-mouse CD31 antibody (BD Pharmingen; 550274, 1:100) at 4°C overnight. The detection was performed as described above. Following nuclear counterstaining, the percentage of Ki-67 positive tumor cells and the microvessel densities (vessels/mm2) were calculated.

***Western Blot***

Western blots were performed as previously described [6]. Antibodies used for western blot included CDK4 (Cell signaling; #12790), CDK6 (Cell signaling; #13331), Rb (Cell signaling; 9309), E2F1 (Cell signaling; #3742), Cyclin D1 (Cell signaling; 2922), FOXM1 (Cell signaling; #20495), α-tubulin (Sigma), βactin (Sigma), GLI1 (Cell signaling; #2643), and the rabbit polyclonal tGLI1 antibody that was developed and validated by our laboratory (9-12)

***Neurosphere Assay***

G48 isogenic cell lines were seeded at a concentration of 4,000 cells/well in 24-well ultra-low attachment plates (Corning #3473) with serum-free Neurobasal Medium (Gibco #21103049) supplemented with 1X GlutaMAX Supplement (Gibco #35050079), 1X B27 (Gibco #17504044), FGF (1ng/mL, Sigma-Aldrich #F0291), EGF (10ng/mL, Sigma-

Aldrich E9644), Sonic Hedgehog Peptide (100ng/mL, Millipore Sigma #S0191) and 1% PenicillinStreptomycin solution (P/S, Corning 30-002-CI). GSC28 isogenic cell lines were seeded at a concentration of 300 cells/well in 24-well ultra-low attachment plates (Corning #3473) with GSC media as described above. Cells were incubated for 24 hours prior to be treated by respective reagents. Fresh media containing treatment reagents were added every 2- 3 days. Neurospheres were counted after 5-9 days culturing under 5X objective. Values 12 were normalized to vehicle control and combination Index (CI) were calculated by the Chou and Talalay method (CompuSyn) (13).
