## Supplementary Figures for "Targeting tGLI1, a novel mediator of tumor therapeutic resistance, using Ketoconazole sensitizes glioblastoma to CDK4/6 therapy and chemoradiation"

### SUPPLEMENTAL FIGURES

#### Overall Survival

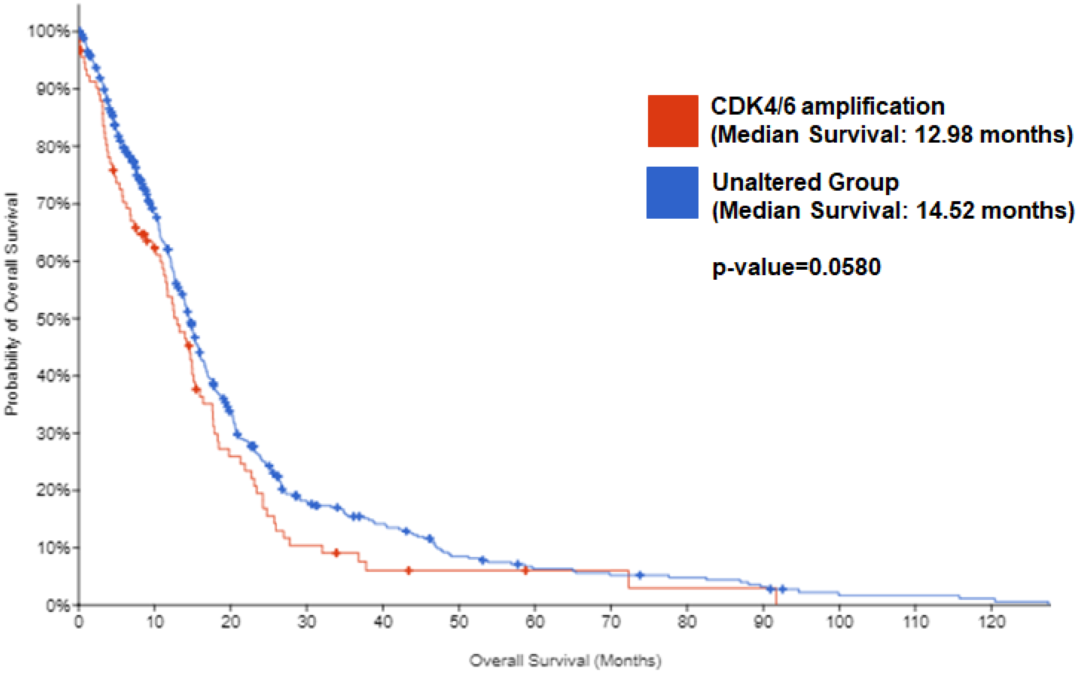


| Number at risk (n) | | | | | | | | | | | | | |
|---|---|---|---|---|---|---|---|---|---|---|---|---|---|
| Altered group | 94 | 52 | 20 | 8 | 4 | 3 | 2 | 2 | 1 | 1 | 0 | 0 | 0 |
| Unaltered group | 501 | 299 | 134 | 63 | 44 | 25 | 17 | 14 | 12 | 8 | 3 | 3 | 2 |

**Supplementary Figure 1:** Patients with CDK4/6 amplification trend toward worse ovetall survival (OS), but it is not significant. Kaplan-Meier analysis of GBM (IDH-WT) patients' OS with or without CDK4/6 amplification from cBioPortal GBM datasets (N=642).

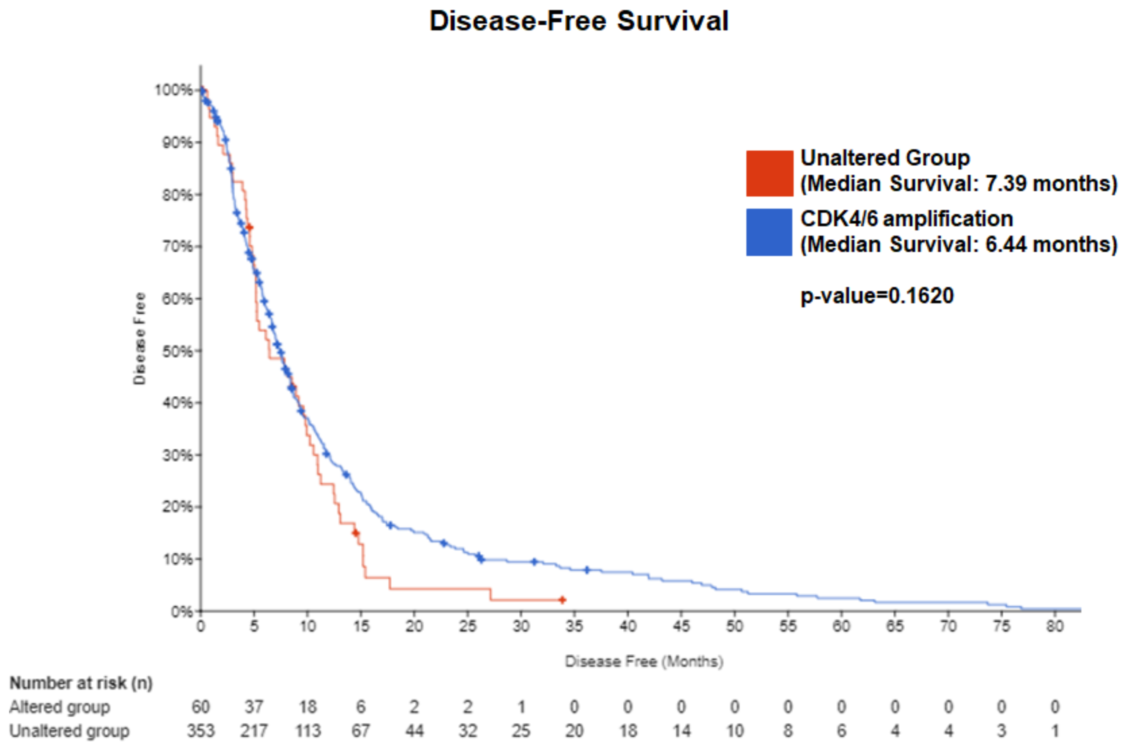


**Supplementary Figure 2: Patients with CDK4/6 amplification trend toward worse disease-free survival (DFS), but it is not significant.** Kaplan-Meier analysis of GBM (IDH-WT) patients' DFS with or without CDK4/6 amplification from cBioPortal GBM datasets. (N=642)

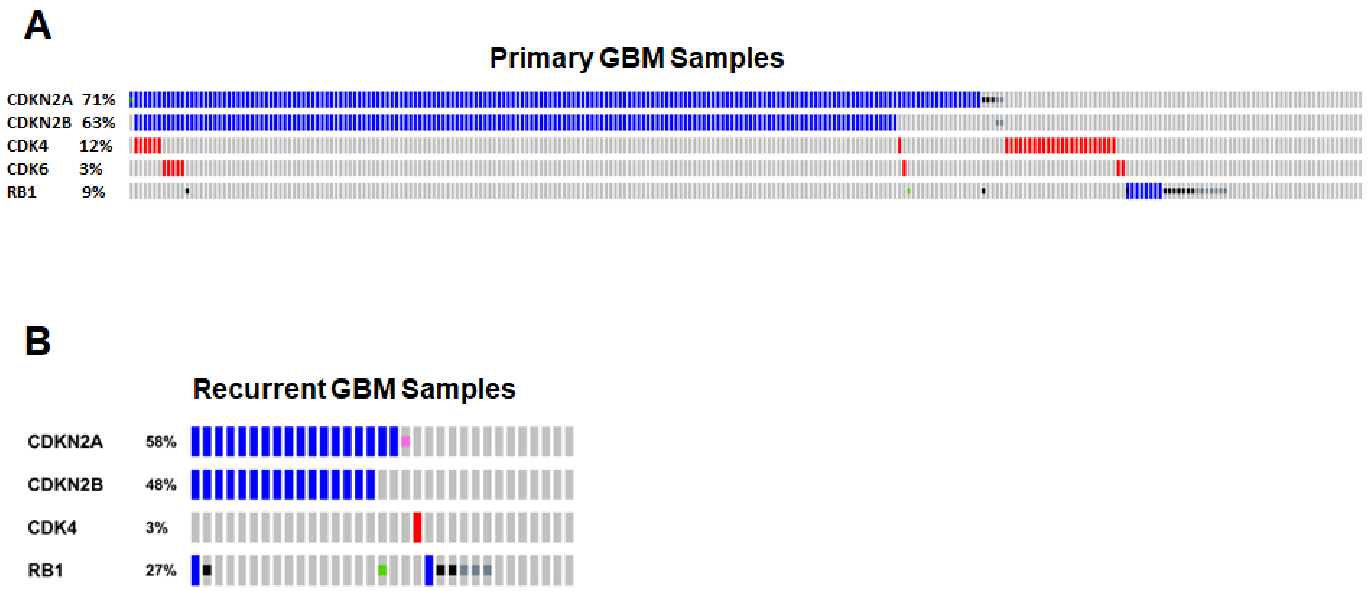


**Supplementary Figure 3: CDK4/6 pathway alterations are frequent in UTHealth Houston GBM patient samples.** (A) Oncoprint of UT Health patient samples CDK4/6 pathway genetic alterations in all patients (N=281). (B) Oncoprint of UT Health patient samples CDK4/6 pathway genetic alterations in recurrent patients (N=33).

A

| Frequency of CDK4 and CDK6 amplification in GBM samples from patients at UTHealth Houston (N=281) | | |
|---|---|---|
| **Genes** | **Newly Diagnosed (N=265)** | **Recurrent (N=33)** |
| **CDKN2A** | 71% | 58% |
| **CDKN2B** | 63% | 48% |
| **CDK4** | 12% | 3% |
| **CDK6** | 3% | - |
| **RB1** | 9% | 27% |

B

| Frequency of CDK4/6 pathway alterations in Matched Primary vs Recurrence (N=33) | | | |
|---|---|---|---|
| **Genes** | **Primary** | **Recurrent** | **Gained Mutations** |
| **CDKN2A** | 76% | 65% | 6% |
| **CDKN2B** | 65% | 59% | 6% |
| **RB1** | 29% | 29% | - |

**Supplementary Figure 4: CDK4/6 pathway alterations are frequent in UTHealth GBM patient samples.**
(A) CDK4/6 pathway alterations are frequent in newly diagnosed (N=265) and recurrent (N=33) GBM patient samples. (B) Frequency of CDKN2A, CDKN2B and RB1 alterations in matched patient samples (N=33).

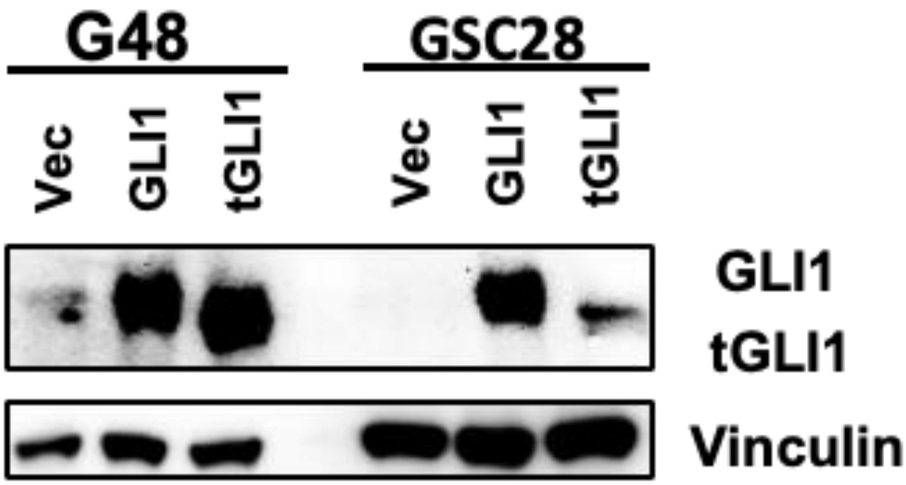


**Supplementary Figure 5: Western Blots demonstrating successful generation of the isogenic G48 and GSC-28 vector, -GLI1, and -tGLI1 cell lines.**

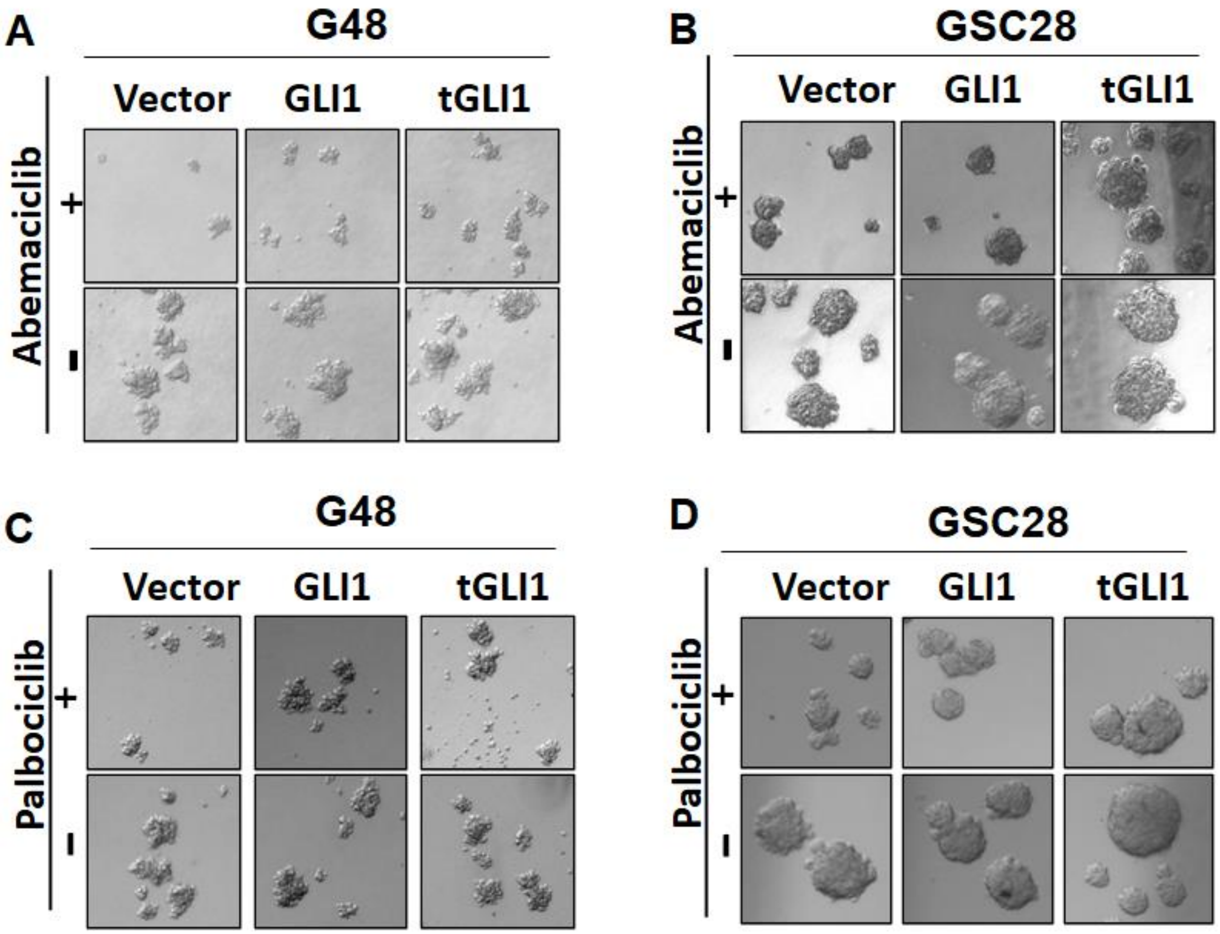


**Supplementary Figure 6: tGLI1 overexpressing GSCs are more resistant to CDK4/6 inhibitor treatment.** (A-D) Representative neurosphere images for isogenic G48 cells treated with 1μM Abemaciclib (A) or 1μM Palbociclib (C) and isogenic GSC-28 cells treated with 5μM Abemaciclib (B) or 5μM Palbociclib (D).

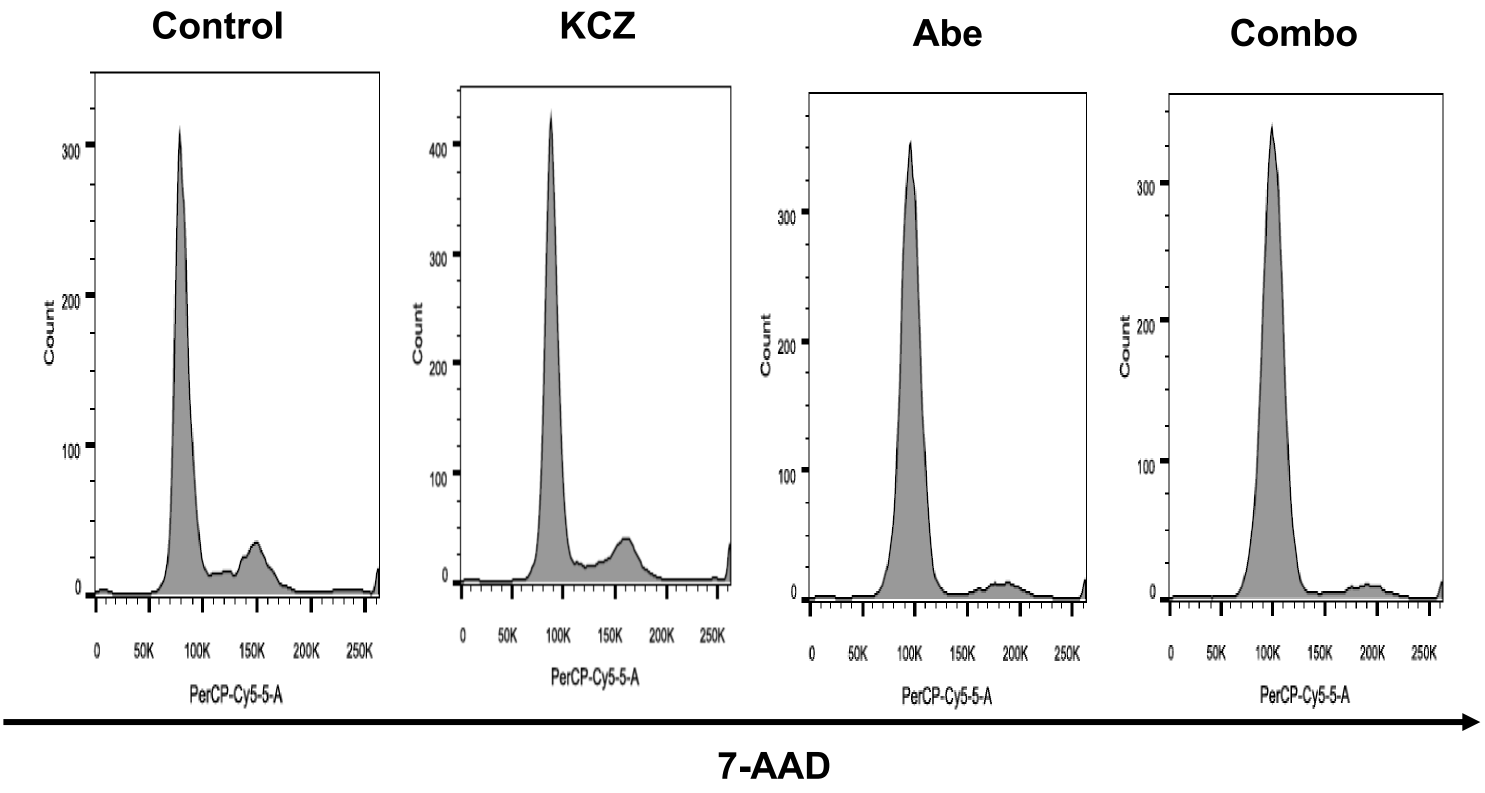


**Supplementary Figure 7: KCZ+Abemaciclib combination therapy significantly decreases GBM proliferation and promotes apoptosis *in vitro*.** Representative histograms of G48-tGLI1 in each phase after treatment with vehicle, KCZ (5uM), Abemaciclib (1uM), or KCZ+Abemaciclib combination for 24 hours

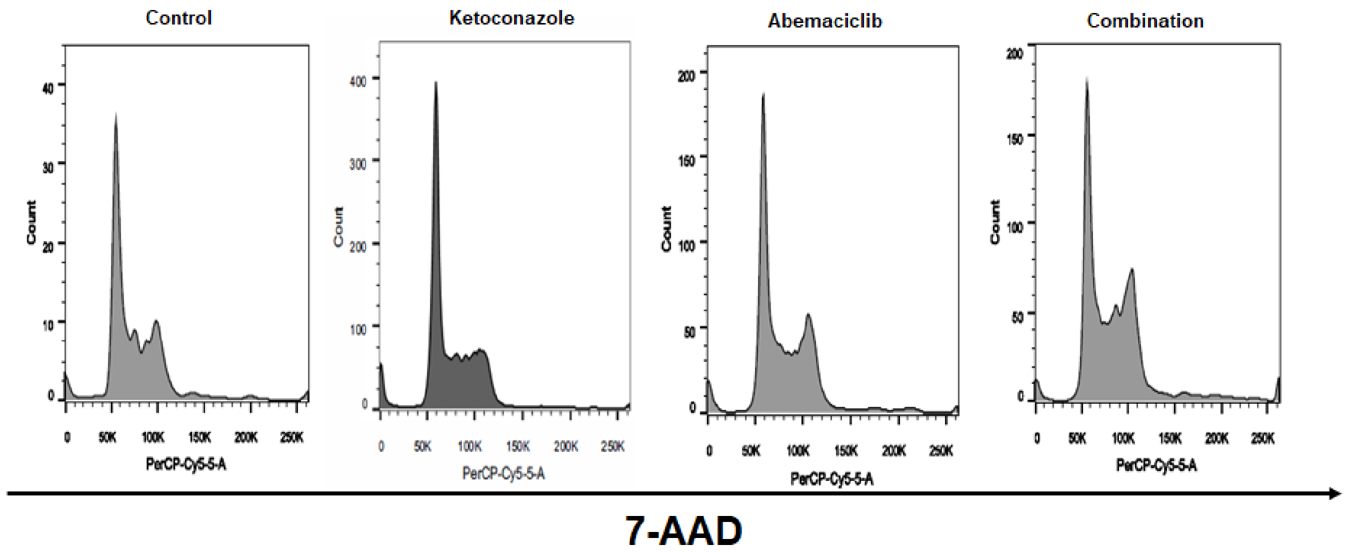


**Supplementary Figure 8: KCZ+Abemaciclib combination therapy significantly decreases GBM proliferation and promotes apoptosis *in vitro*.** Representative histograms of GSC28-tGLI1 in each phase after treatment with vehicle, KCZ (5uM), Abemaciclib (1uM), or KCZ+Abemaciclib combination for 24 hours.

| Total flux (photon/sec) | |
|---|---|
| Control | 2.85E+07 |
| KCZ | 1.17E+07 |
| Abe | 1.53E+07 |
| Both | 4.69E+06 |

**Supplementary Figure 9: KCZ+Abemaciclib combination therapy significantly decreases tumor growth *in vivo.*** Table above shows the quantitative total flux from IVIS images at 27 days after intracranial inoculation (in Figure 6B). Combination treatment has the lowest flux compared to all other groups.

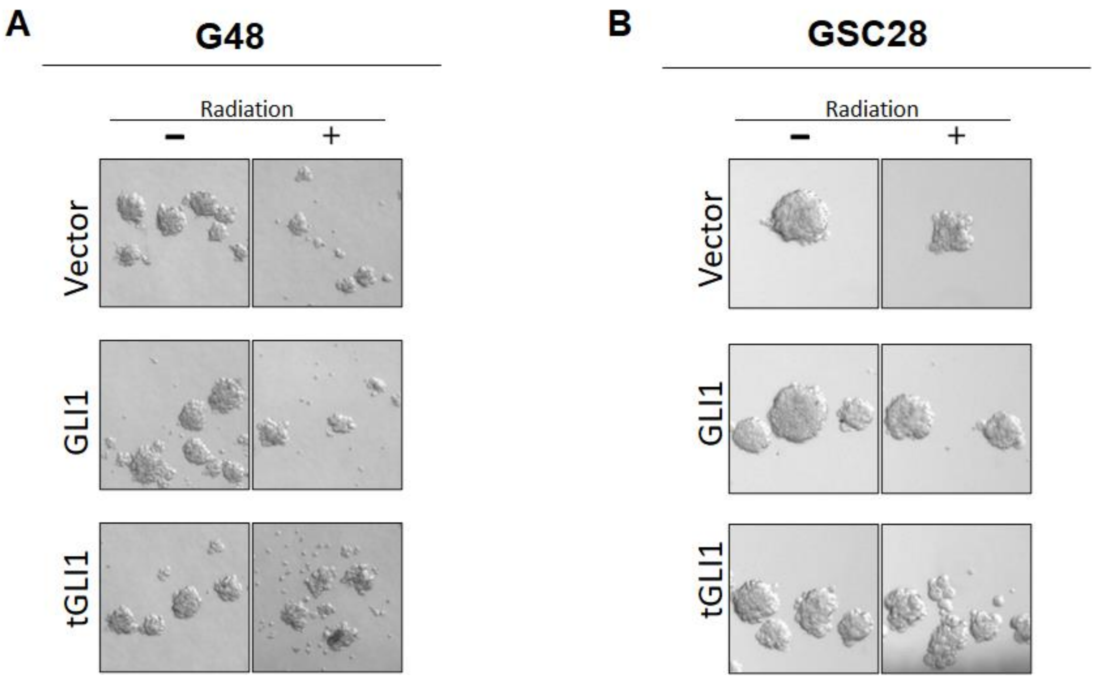


**Supplementary Figure 10: tGLI1 overexpressing GSCs are more resistant to radiation treatment. (**A-B) Representative neurosphere images for isogenic G48 and GSC-28 vector, -GLI1, and -tGLI1 cells treated with radiation treatment (5Gy and 2Gy respectively).
