## Supplementary material for "Targeting tGLI1, a novel mediator of tumor therapeutic resistance, using Ketoconazole sensitizes glioblastoma to CDK4/6 therapy and chemoradiation": Original Western Blots

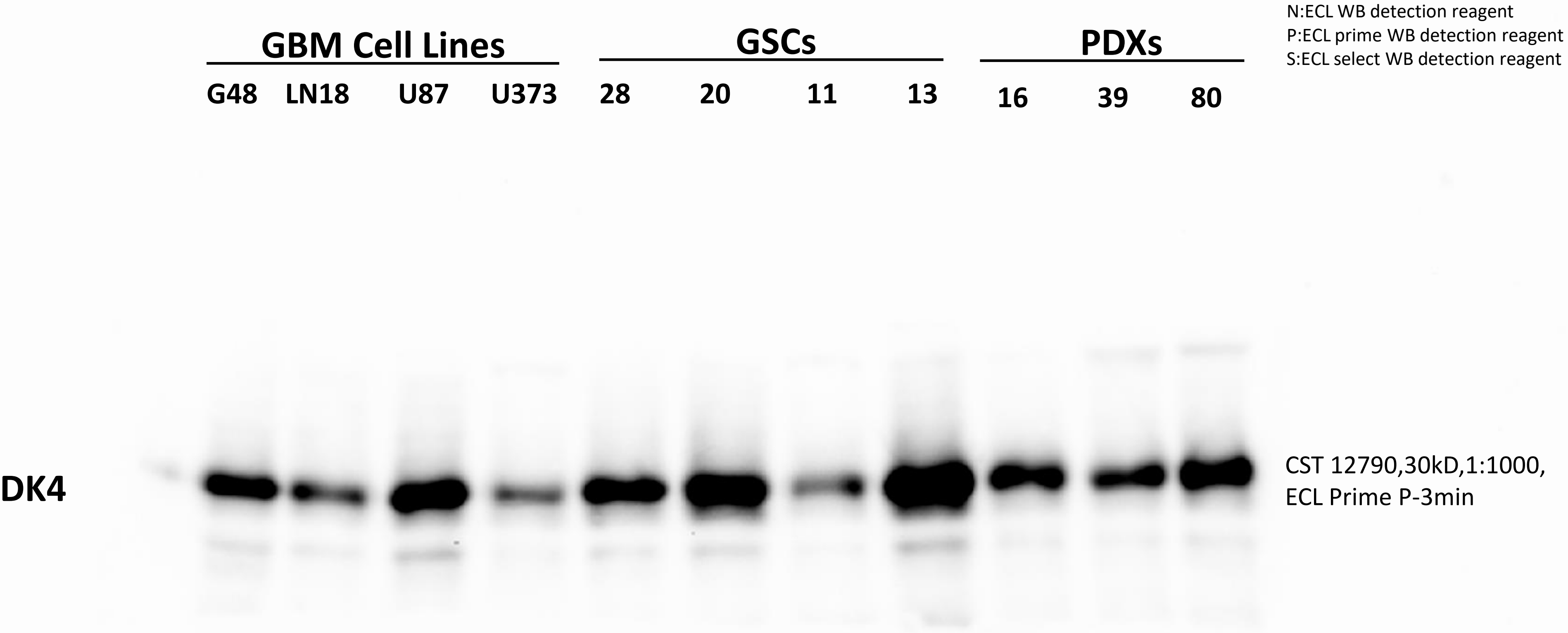


Supplemental Material 1: original blot for CDK4 Western blot for Figure 2J

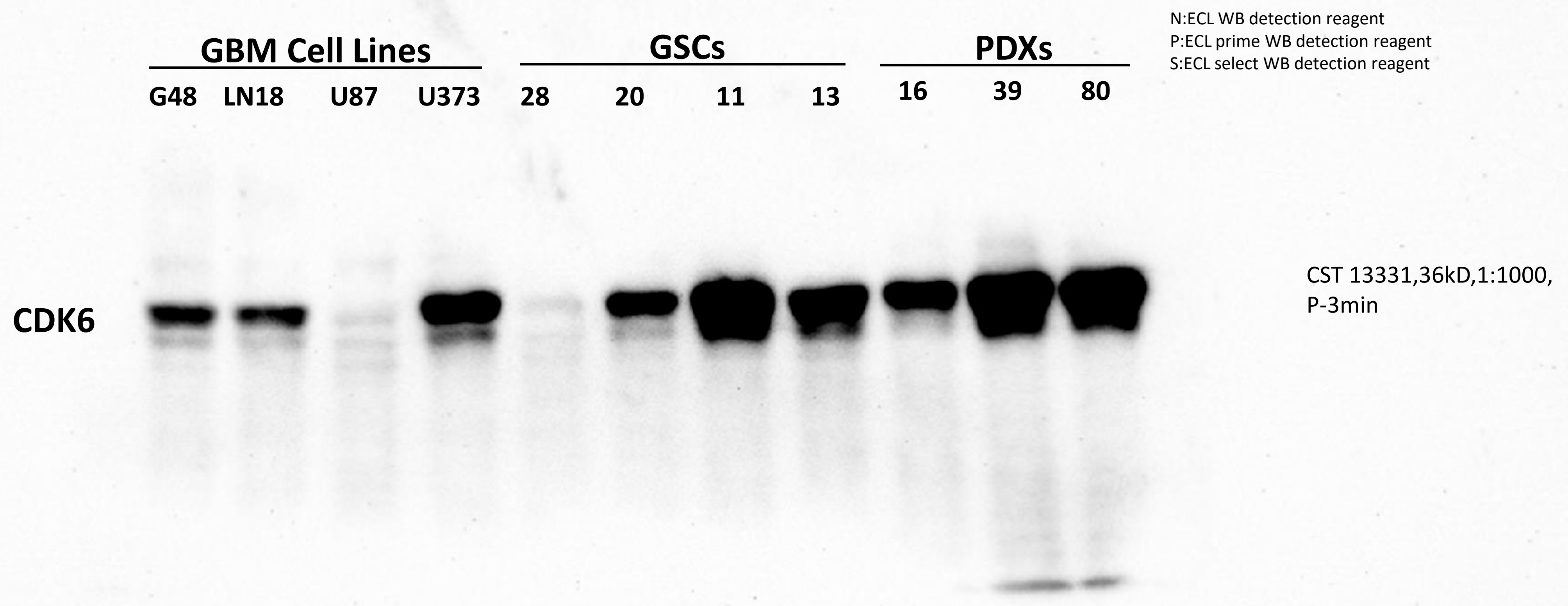


Supplemental Material 2: original blot for CDK6 Western blot for Figure 2J

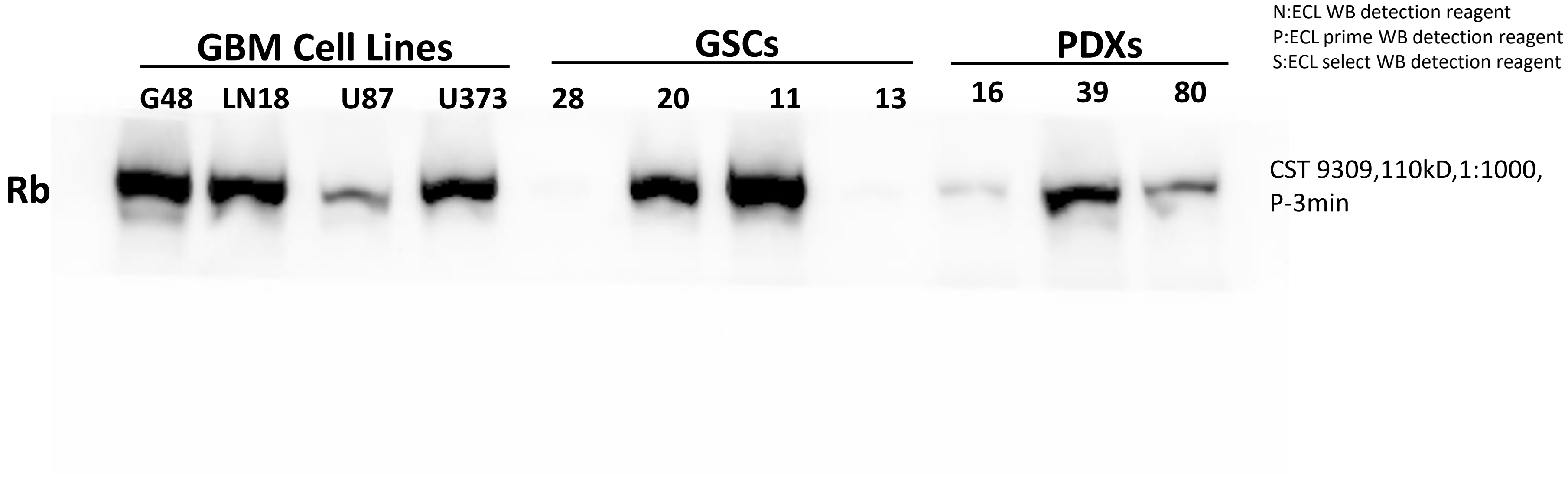


Supplemental Material 3: original blot for Rb Western blot for Figure 2J

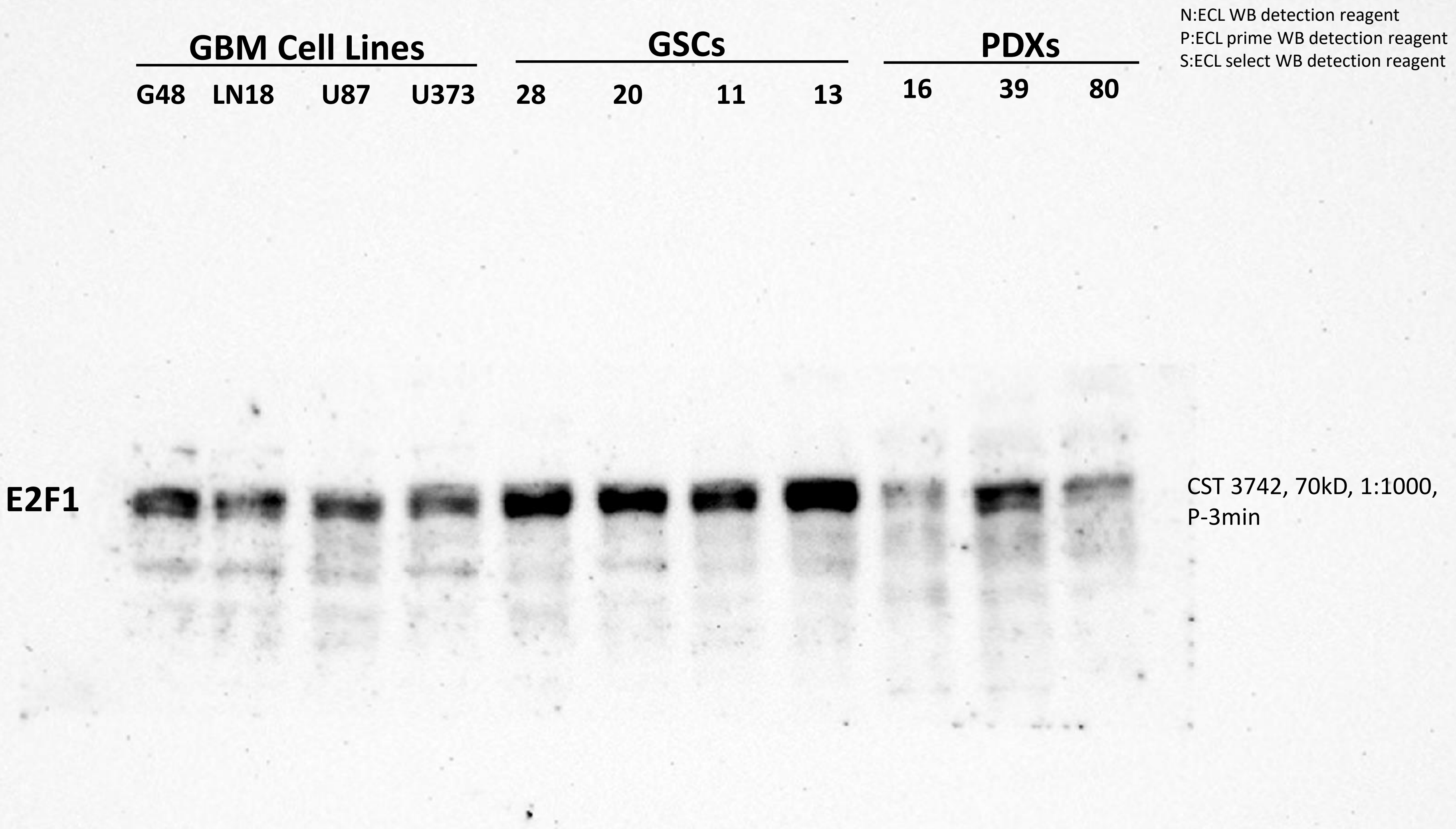


Supplemental Material 4: original blot for E2F! Western blot for Figure 2J

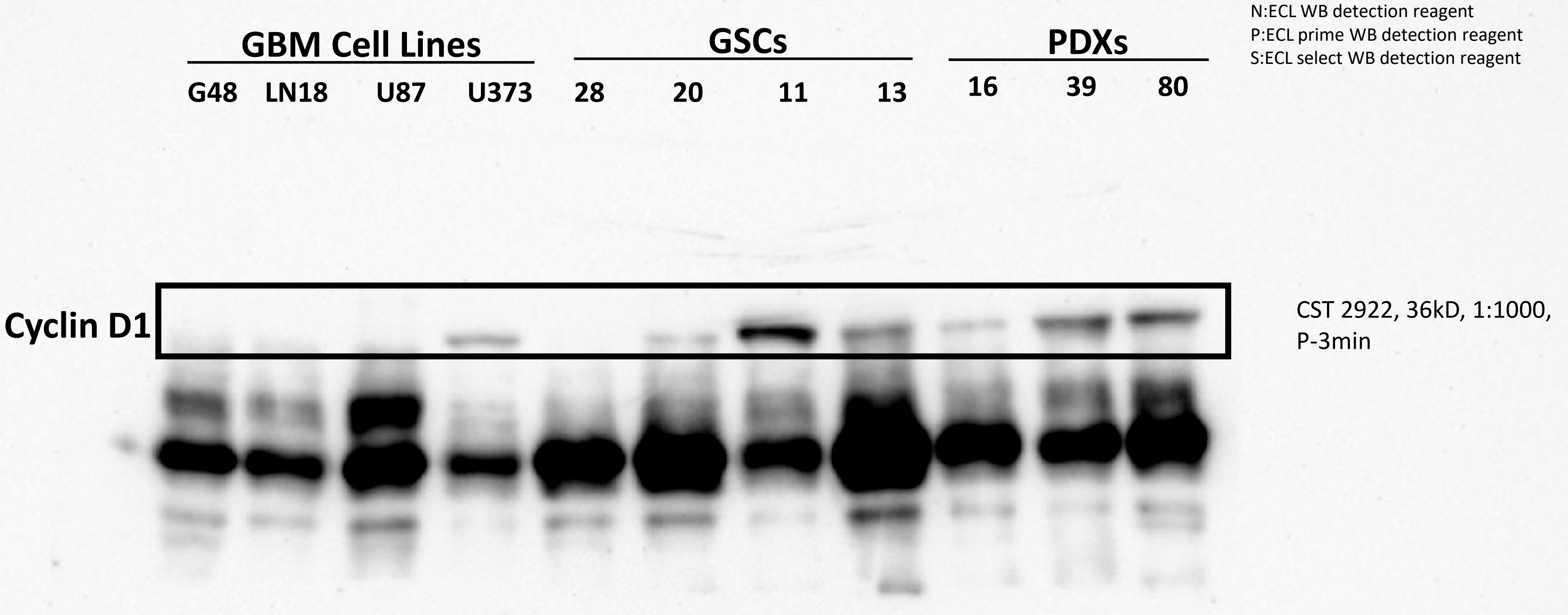


Supplemental Material 5: original blot for Cyclin D1 Western blot for Figure 2J

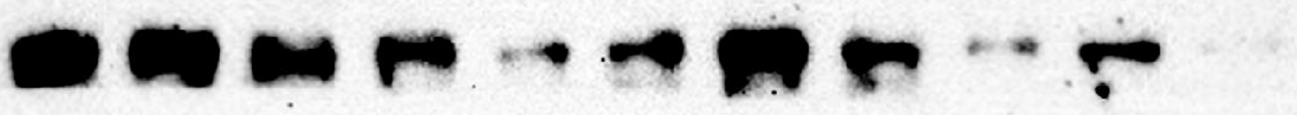


Supplemental Material 6: original blot for FOXM1 Western blot for Figure 2J

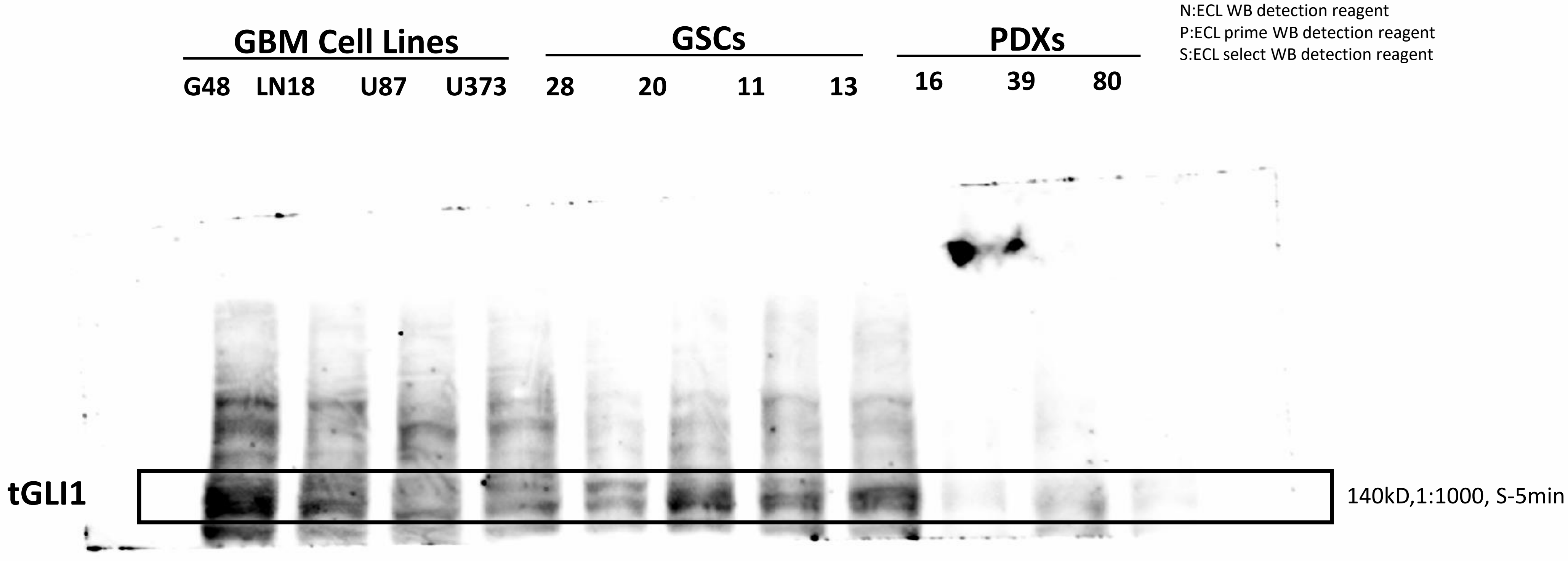


Supplemental Material 7: original blot for tGLI1 Western blot for Figure 2J

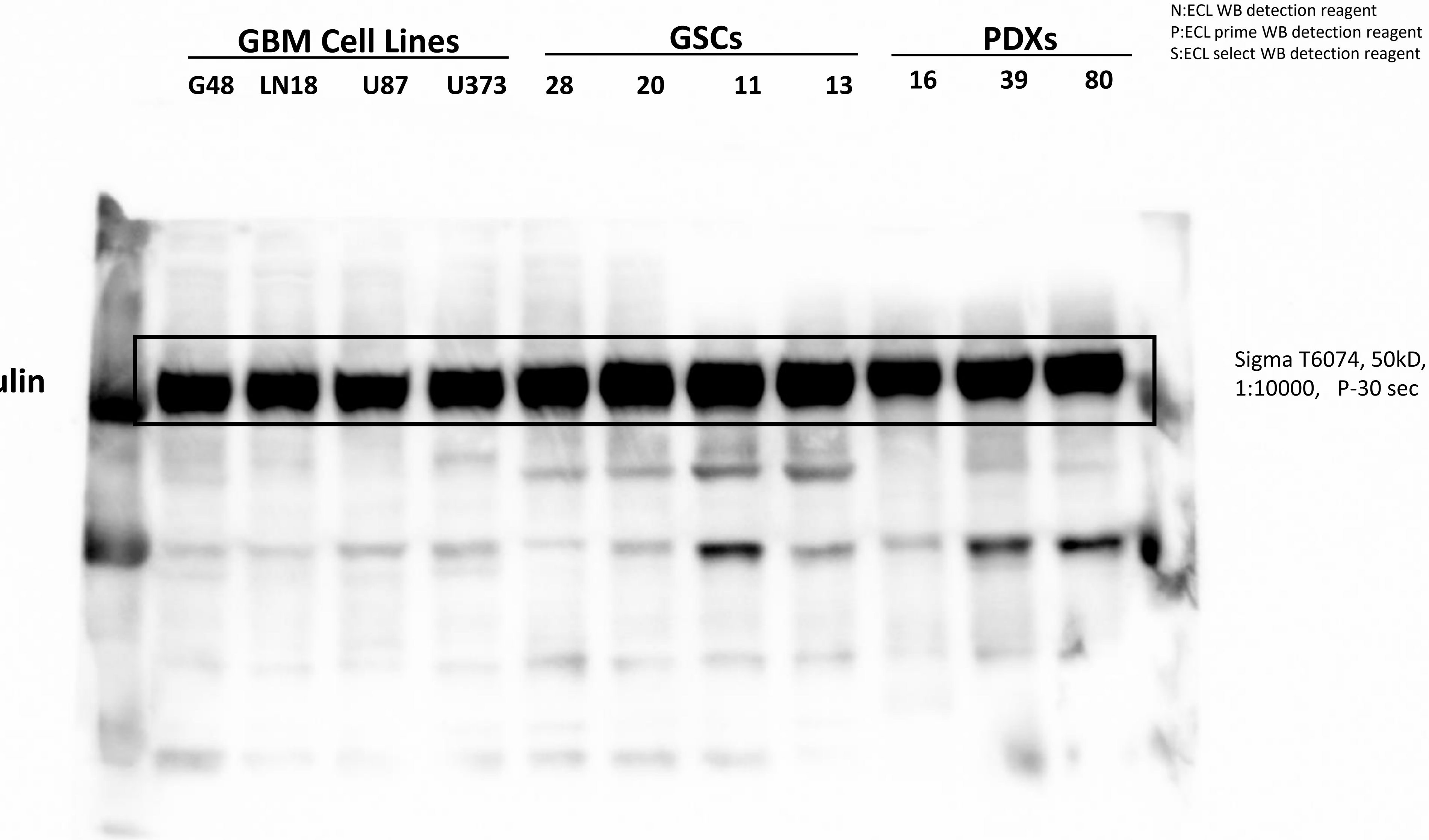


Supplemental Material 8: original blot for CDK6 Western blot for Figure 2J

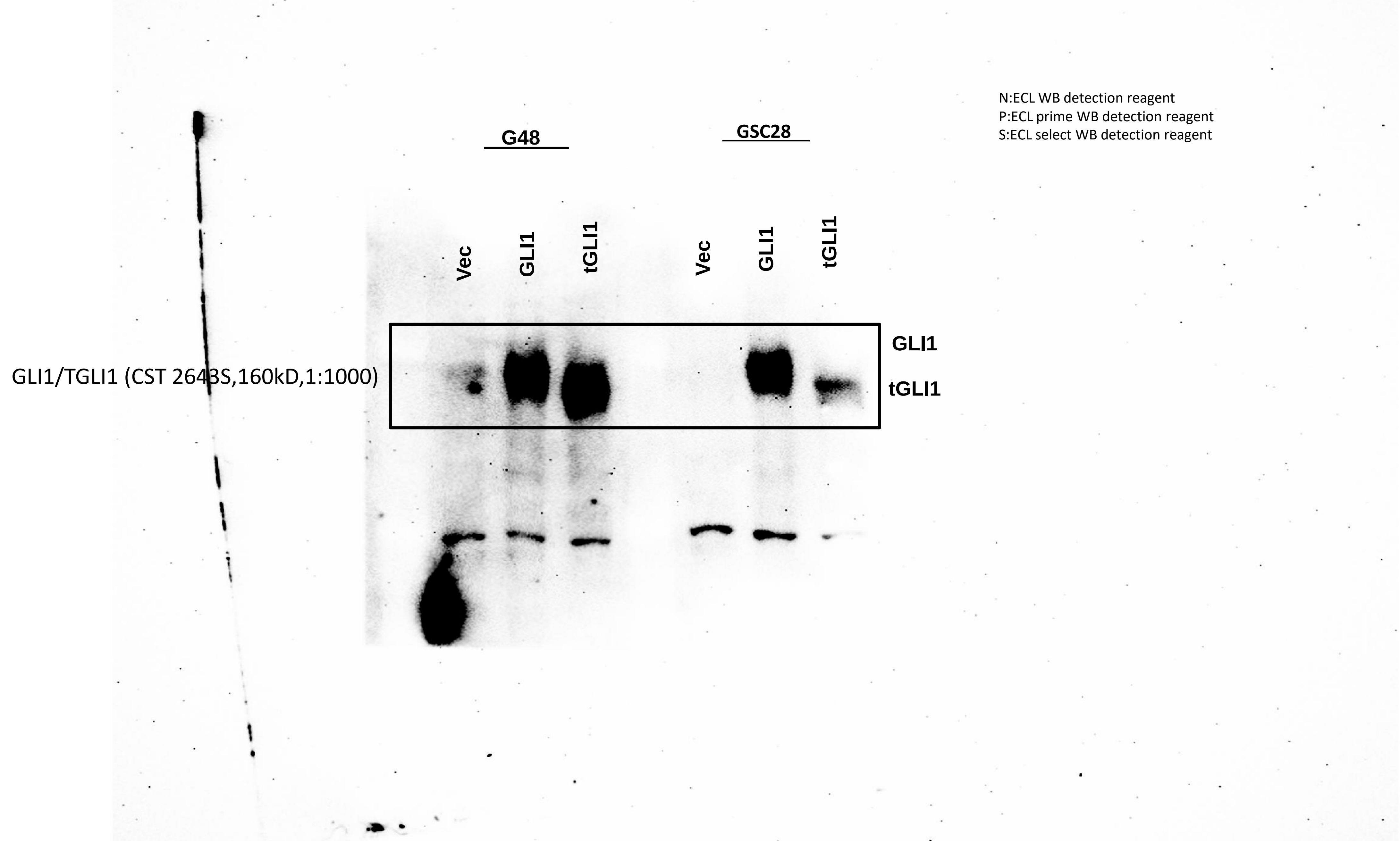


Supplemental Material 9: original blot for GLI1/tGLI1 Western blot for Figure 3A

N:ECL WB detection reagent
P:ECL prime WB detection reagent
S:ECL select WB detection reagent

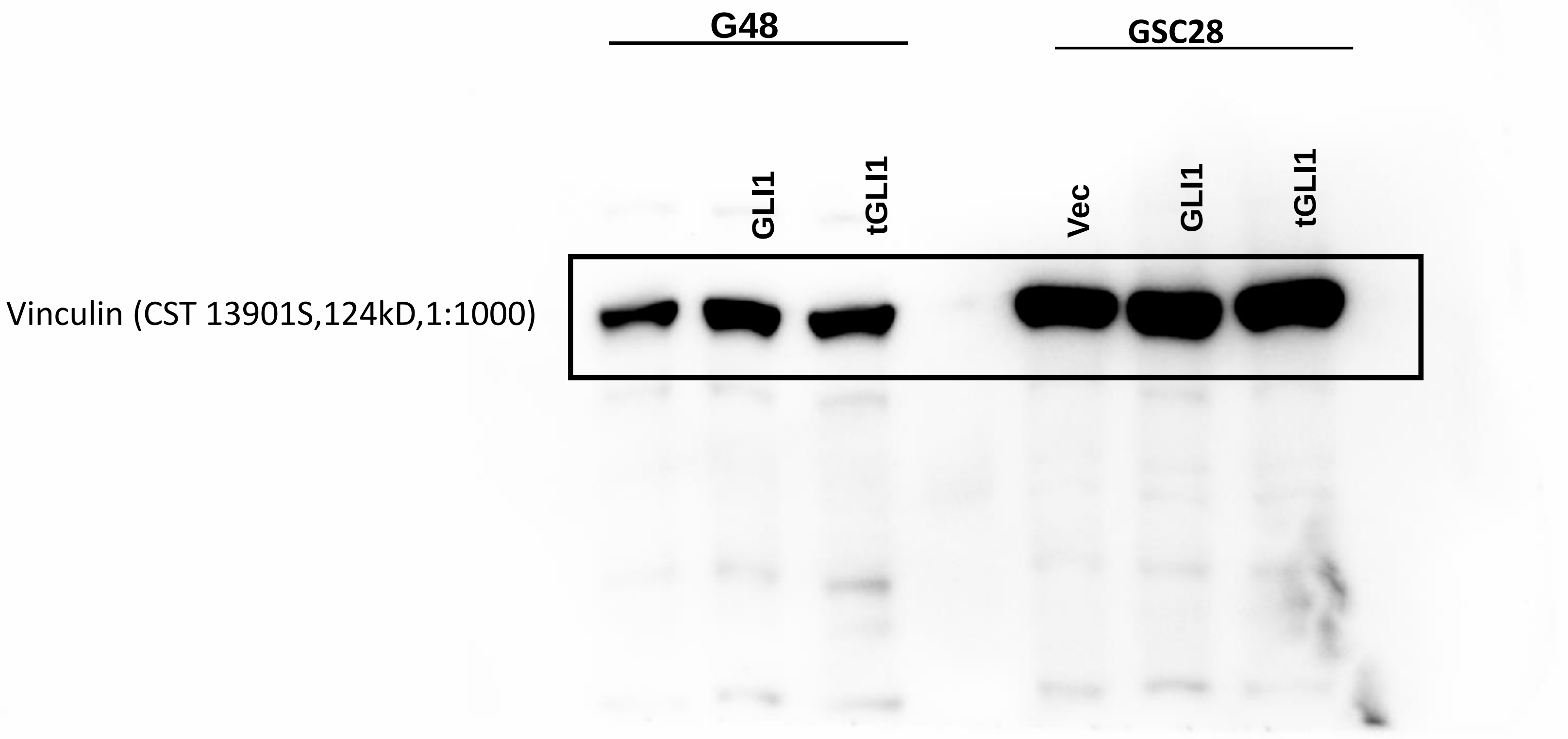


Supplemental Material 10: Original blot for Vinculin Western blot for Figure 3A
